# Estrogen receptor alpha mutations regulate gene expression and cell growth in breast cancer through microRNAs

**DOI:** 10.1101/2022.10.07.511340

**Authors:** Spencer Arnesen, Jacob T. Polaski, Zannel Blanchard, Kyle S. Osborne, Alana L. Welm, Ryan M. O’Connell, Jason Gertz

## Abstract

Estrogen receptor α (ER) mutations occur in up to 30% of metastatic ER-positive breast cancers. Recent data has shown that ER mutations impact the expression of thousands of genes not typically regulated by wildtype ER. While the majority of these altered genes can be explained by constant activity of mutant ER or genomic changes such as altered ER binding and chromatin accessibility, as much as 33% remain unexplained, indicating the potential for post-transcriptional effects. Here we explored the role of microRNAs in mutant ER-driven gene regulation and identified several microRNAs that are dysregulated in ER mutant cells. These differentially regulated microRNAs target a significant portion of mutant-specific genes involved in key cellular processes. When the activity of microRNAs is altered using mimics or inhibitors, significant changes are observed in gene expression and cellular proliferation related to mutant ER. An in-depth evaluation of miR-301b led us to discover an important role for *PRKD3* in the proliferation of ER mutant cells. Our findings show that microRNAs contribute to mutant ER gene regulation and cellular effects in breast cancer cells.

## Introduction

Estrogen receptor α (ER) is a steroid hormone receptor that is activated by estrogens and is expressed in 70% of breast cancers. ER plays an important role in driving the progression of these tumors by regulating genes to promote growth and proliferation. Due to the availability of effective anti-estrogen therapies, including selective estrogen receptor modulators and aromatase inhibitors (AI), ER-positive breast cancers generally have a good prognosis with a five-year survival rate of greater than 90% (1). However, many ER-positive breast cancer patients eventually develop resistance to hormone therapies and experience recurrent metastatic disease (2,3). Activating mutations in the ligand binding domain (LBD) of ER have been observed in approximately 30% of hormone therapy-resistant ER-positive breast cancers (4–6). Over the last decade, several studies have improved our understanding of ER mutations and the mechanisms by which these mutations lead to breast cancer recurrence (4–18). Structural changes in ER’s LBD as a result of ER LBD mutations confer an active conformation and ligand-independent activity to unliganded mutant ER, which drives large transcriptional changes in the absence of estrogens (5,7,9,11–16). Several studies have shown that this dramatic change in gene expression can drive increased cellular proliferation, invasion, and metastasis in breast cancer cells expressing mutant ER (4,5,9,13,14,16–18). Recently, we showed that over half of these gene expression changes are driven directly by the constitutive activity of ER (16). Additionally, changes in ER’s genomic binding and widespread alterations to chromatin accessibility, accompanied by binding of other transcription-regulating factors such as CTCF, FOXA1, and OCT1, can explain an additional 20-30% of transcriptional changes observed in ER mutant breast cancer cells. However, the intermediate factors driving these transcriptional changes and the causes of the remaining unexplained gene expression changes are unknown.

Since ER has been shown to activate several microRNAs (19,20), we hypothesized that regulation of microRNAs(miRNAs) could contrubte to altered gene expression in ER mutant cells. miRNAs are short RNA molecules with an average length of 22bps that are transcribed by RNA polymerase II. miRNAs typically bind directly to the 3’ untranslated region (UTR) of mature messenger RNAs (mRNAs), which usually leads to decreased translation and increased degradation of the target mRNAs (21,22). Many miRNAs have been described to have either oncogenic or tumor-suppressive roles, and the altered expression of these miRNAs often contributes to cancer-associated cellular processes, including increased proliferation, migration, and metastasis (23–26). These effects are often produced when miRNAs alter the activity of important signaling pathways in cancer cells (27). Altered activity of these pathways by miRNAs could provide therapeutic opportunities. In addition, miRNA mimics and inhibitors have recently been considered as a potential therapeutic approach for treating some diseases (26,28–30). The possible use of miRNA mimics and inhibitors as cancer treatments in the future makes understanding the role of miRNAs in cancer an important subject.

Activated ER homodimerizes and binds to genomic elements in the genome, termed estrogen response elements (ERE), where it recruits cofactors that drive the transcription of nearby genes via RNA Polymerase II. Because miRNAs are transcribed by RNA Polymerase II, they could be directly regulated by ER or by other transcription factors downstream of ER signaling. Indeed, some miRNAs have previously been shown to be regulated by ER activation (19,20). These ER regulated miRNAs, along with many other miRNAs, have been shown to contribute to cancer progression in ER-positive breast cancers (27,31–38). Despite the clear role of miRNAs in contributing to cancer progression both broadly and in the specific case of ER-positive breast cancers, the role of miRNAs in recurrent mutant ER-positive breast cancers has thus far remained unexplored. The potential for ER to directly or indirectly regulate miRNA transcription suggests that differential expression of miRNAs could occur in ER mutant cells and could contribute to the large transcriptional changes we and others have observed in mutant ER-positive breast cancer cells.

Here, we investigate the role of miRNAs in regulating gene expression in breast cancer cell lines we developed previously, which express wildtype (WT) ER or heterozygously express either the Y537S or the D538G mutation, the two most common ER LBD mutations, from the endogenous ER locus (16). We identified many miRNAs that are differentially expressed in ER mutant cells compared to WT ER cells. We examined the effects of several of these miRNAs, including miR-301b, miR-181c, miR-331, and let-7f, on gene expression and found that they directly or indirectly alter the expression of hundreds of genes previously identified as being differentially expressed in ER mutant cells compared to WT cells (mutant-specific genes) (16). These miRNAs also impact the ability of ER mutant cells to proliferate, a process that is significantly increased in ER mutant cells compared to their WT counterparts. In particular, we identify miR-301b as being significantly down-regulated in ER mutant cells, which led to altered expression of mutant-specific genes and increased proliferation. Decreased miR-301b expression increases the RNA expression of *PRKD3*, a kinase regulator of several cellular signaling pathways, including the RAS/MEK/ERK pathway (39–42). We found that inhibiting PKD3 protein leads to significantly decreased proliferation in ER mutant cells. In summary, we discovered that several miRNAs, including miR-301b, play an important role in driving ER mutant gene regulation and contribute to increased cellular proliferation in ER mutant breast cancer cells. We show that these miRNAs and their targeted pathways could provide alternative therapeutic strategies for treating mutant ER-positive breast cancer.

## Materials and Methods

### Cell culture

The generation of T-47D ER mutant and WT clones was described in Arnesen et al (16). Cells were cultured in RPMI1640 Media (ThermoFisher Scientific) supplemented with 10% FBS (Sigma-Aldrich) and 1% penicillin-streptomycin (ThermoFisher Scientific). Five days prior to all experiments, except those otherwise specified, cells were moved to growth in hormone-deprived media which was composed of phenol red-free RPMI1640 Media (ThermoFisher Scientific) supplemented with 10% charcoal-stripped FBS (ThermoFisher Scientific) and 1% penicillin-streptomycin. Prolonged 25 day E2 treatment of WT cells was performed as described in Arnesen et al (16).

### miRNA expression analysis

ER mutant or WT T-47D cells were plated in 100mm dishes and grown in estrogen-deprived media for five days prior to treatment with DMSO (0.1%) for 8 hours or for *ESR1* knock-down experiments, cells were treated with DMSO (0.1%) and transfected with siRNAs targeting *ESR1* or a non-targeting siRNA control, as described below, for 48 hours prior to RNA harvest. Cells were washed with phosphate-buffered saline (PBS; ThermoFisher Scientific) and treated with RLT plus (Qiagen) lysis buffer with 1% beta-mercaptoethanol (BME; Sigma-Aldrich) for cell lysis. Cells were passed through a 21-gauge needle (Sigma-Aldrich) to facilitate genomic DNA lysis. For PDX lines, tissue samples were used from seven PDX lines (WT: HCI-003 and HCI-011; L536P: HCI-005, HCI-006, HCI-007; Y537S: HCI-013, HCI-013EI). PDX samples were surgically removed from mice and flash frozen. PDX samples were not treated. Approximately 20mg of tissue per sample was homogenized in RLT plus lysis buffer with 1% beta-mercaptoethanol using gentleMACS M-tubes and a gentleMACS Octo Dissociator (Miltenyi Biotec). RNA extraction for both T-47D and PDX samples was performed using a Quick-RNA RNA Miniprep kit (Zymo Research), and RNA was measured using a Qubit fluorometer. For each sample, 200ng of total RNA was submitted for miRNA expression analysis using NanoString’s nCounter Human v3 miRNA expression assay. NanoString miRNA expression assay results in the form of raw counts were analyzed using NanoString’s nSolver software. Raw counts were normalized using two approaches. The first approach used the geometric mean for each sample of the top 100 highest expressed miRNAs across all samples to determine a normalization factor for each sample. The second approach used the geometric mean for each sample of the five spike-in miRNAs profiled for the assay to generate the normalization factors. Normalized counts were then log-scaled and used to perform *t* tests to identify significantly differentially expressed miRNAs. Because this was a hypothesis generating step, we considered differentially expressed miRNAs from both normalization approaches in our study. A paired *t* test was used for calling differentially expressed miRNAs from the *ESR1* knock-down experiment due to clonal variability between clones of the same genotype. The DIANA TarBase database (43) was used to compare putative target genes of differentially expressed miRNAs with mutant-specific genes described in Arnesen et al (16). Significant enrichments were defined using a hypergeometric test as implemented in R using phyper.

### Cell transfections

After cells were grown in hormone-deprived media for five days, cells were transfected with a miRNA mimic, miRNA inhibitor, or an siRNA. mirVana miRNA mimics and inhibitors were used for 15 miRNAs (ThermoFisher Scientific). Mimics and inhibitors were transfected at a 10nM final concentration using the RNAiMAX transfection reagent (ThermoFisher Scientific) per the manufacturer’s instructions. siRNAs against *PRKD3* (Integrated DNA Technologies’ TriFECTa RNAi kit) or against *ESR1* (Horizon siGENOME SMARTPool siRNA reagents, M-003401-04-0005) were used to knock-down *PRKD3* or *ESR1* transcript levels, respectively (sup. table S4). To test the efficacy of these siRNAs, WT cells were transfected with 0.1nM, 1nM, or 10nM concentrations of three anti-*PRKD3* or the pooled anti-*ESR1* siRNAs using the RNAiMAX transfection reagent. The two siRNAs achieving the best knock-down of *PRKD3* or the pooled anti-*ESR1* siRNAs were used at a 10nM concentration in subsequent siRNA transfection experiments.

### Proliferation assays

Cells were plated in 100mm dishes and grown in hormone-deprived media for four days prior to plating in 96-well plates at approximately 12,500 cells per well and allowed to adhere for another day in hormone-deprived media. 24 hours after 96-well plating, cells were treated with 1% DMSO and a miRNA mimic/inhibitor, siRNA, or drug, depending on the experiment. Cells were then placed on the IncuCyte Zoom Live Cell Imaging Platform (Sartorius) and proliferation was monitored for 48 hours with 10x magnification images obtained at 2-hour intervals. Confluence was measured for three or four replicates, depending on the experiment, and percent confluence values were normalized to the initial confluence and then log2 transformed. Linear regression analysis using a general linear Wald test, as implemented by glm() in R, was used to compare the slopes of transformed confluence values over time and to determine significant differences in slopes.

### RNA-seq

T-47D cells were grown in 100mm plates in hormone-deprived media for four days then plated into 12-well plates at approximately 50% confluence and allowed to adhere for 24 hours. Cells were then transfected with a miRNA mimic or inhibitor for a given miRNA or with a mimic or inhibitor negative control at a 10nM final concentration. 24 hours post-transfection, cells were collected, washed with PBS, then treated with RLT Plus lysis buffer supplemented with 1% BME. RNA was extracted using a Quick-RNA RNA Miniprep kit (Zymo Research), and RNA was measured using a Qubit fluorometer. Poly-A selected RNA-seq libraries were generated using a KAPA Stranded mRNA-Seq kit (KAPA Biosystems) and 500ng RNA per sample as starting material. Libraries were sequenced using the Illumina NovaSeq 6000 platform. The Hisat2 spliced aligner was used to align the resulting fastq files to the hg19 human genome build (44). SAM files were then converted to BAM files using SAMtools (45). Genes included in the University of California Santa Cruz’s (UCSC) Known Genes table were assigned counts using the Subread package’s featureCounts program using BAM files as inputs (46). Counts were then normalized and analyzed for differential expression using the DESeq2 package in R (47). Two clones for each ER genotype (Y537S, or D538G) were used for RNA-seq and clones of the same genotype were used as biological duplicates in our differential expression analysis. Significantly up-or down-regulated genes (adjusted p-value <0.05) were identified by comparing results from miRNA mimic or inhibitor treatment to the appropriate negative control treatment in clones of the same ER genotype. Genes that were identified as differentially expressed were compared to mutant-specific genes as defined previously (16). miRNA-altered mutant-specific genes were compared with putative miRNA target genes for the corresponding miRNA using the miRDB microRNA Target Prediction Database (48). Significant enrichments were determined using a hypergeometric test as implemented in R using phyper with a p-value cutoff of <0.05. Gene ontology and pathway analysis was performed by submitting genes on interest to Ma’ayan lab’s Enrichr online tool (49). Adjusted p-values were used to identify significantly enriched ontologies and pathways. Venn diagrams were made using the eulerr package.

### Quantitative PCR

WT T-47D cells were grown in regular media. Cells were lysed using RLT plus (Qiagen) plus 1% beta-mercaptoethanol (Sigma-Aldrich) and total RNA was extracted using a Quick RNA Miniprep kit (Zymo Research). qPCR was performed using a Power SYBR Green RNA-to-CT 1-Step Kit (ThermoFisher Scientific) and 50ng of RNA per sample as input. A CFX Connect Real-Time light cycler (Bio-Rad) was used for thermocycling and light detection. A ΔΔCt method was used to calculate expression levels with TBP used as the control. Duplicates were measured for each sample. Primer sequences for *PRKD3* and *TBP* primers are listed in Supplemental Table S4. Significant changes in relative expression were determined using a one-tailed Student’s *t* test using t.test in R, with a p-value cutoff of <0.05.

### Immunoblot analysis

ER mutant or WT cells were grown in 6-well format. Cells were transfected with an siRNA or a miRNA mimic/inhibitor and then collected and lysed 48 or 72 hours post transfection using RIPA buffer (1x PBS, 1% NP-40, 0.5% sodium deoxycholate, 0.1% SDS) supplemented with protease and phosphatase inhibitor (ThermoFisher Scientific, A32959). Total protein concentration was determined using the Bradford quantification assay (Bio-Rad). For electrophoresis, 20-30 µg of total protein was diluted with 4X NuPAGE^TM^ LDS Sample Buffer (ThermoFisher Scientific, NP0007) and run on a NuPAGE^TM^ 4-12% Bis-Tris polyacrylamide gel (ThermoFisher Scientific, NP0336BOX) at 150-200V. Protein was then transferred to nitrocellulose membrane (LI-COR Biosciences, 926-31092) in transfer buffer (ThermoFisher Scientific, NP00061) supplemented with 20% methanol (Millipore Sigma, 179337) overnight at 4°C with constant stirring. Blots were blocked with Odyssey Blocking Buffer (LI-COR Biosciences, 927-60001) for 1 hour at 4°C and probed with the following antibodies overnight with shaking at 4°C: PRKD3 (Proteintech, 12785-1-AP, 1:1000 dilution), ER (Santa Cruz Biotechnology, sc-543, 1:1000 dilution), Beta-Actin (Santa Cruz Biotechnology, [C4] sc-4778, 1:1000 dilution), and GAPDH (Abcam, [6C5] ab8245, 1:2000 dilution). The blot was then washed three times with 1X TBST buffer at room temperature and probed with 1:10000 dilutions of mouse secondary antibody (LI-COR Biosciences IRDye^®^ 680RD Goat anti-mouse IgG, 926-68072) and/or rabbit secondary antibody (LI-COR Biosceiences IRDye^®^ 800CW Donkey anti-rabbit IgG, 926-32213) for 1 hour at room temperature.Following incubation with secondary antibodies, the blot was washed three times with 1X TBST buffer, visualized using an Azure^TM^ Biosystems 600 imaging system, and band intensity was quantified using Image Studio Lite software (Image Studio Lite Software, LI-COR, Lincoln NE). Statistical analysis of protein abundance was performed using a one-tailed Student’s *t* test using t.test in R, with a p-value cutoff of <0.05.

### Drug treatments

Cells were grown for a single clone for each ER genotype (WT, Y537S, and D538G) in 100mm dishes in hormone-deprived media for 4 days and then plated in 96-well plates at approximately 12,500 cells per well and allowed to adhere in hormone-deprived media for 24 hours. Cells were then treated with varying doses (0.01 uM, 0.1 uM, 0.5 uM, 1 uM, 5 uM, 10 uM, or 100 uM) of the PKD inhibitor CRT0066101 (Selleck Chemicals, S8366) or with a vehicle control (H_2_O) and analyzed for proliferation. Experiments were performed in triplicate. Proliferation rates were calculated as described above and IC_50_ values were calculated using Prism 9 software.

### Statistical analysis and packages

All statistical analyses, except those for gene ontology and pathways analysis, were performed using R version 3.5.2 or 3.5.3.

## Results

### ER mutations alter the expression of miRNAs

Previously, we identified thousands of genes that are differentially expressed in D538G or Y537S ER mutant cells compared to WT ER cells (mutant-specific genes) regardless of the presence or absence of 17β-estradiol (E2). Post-transcriptional mechanisms, such as differential expression of miRNAs in ER mutant cells, may contribute to mutant-specific gene expression. To determine whether there is differential expression of miRNAs in ER mutant cells, we utilized T-47D breast cancer cells, developed previously, which heterozygously express either the D538G or Y537S mutant *ESR1* allele from the endogenous *ESR1* locus (16). ER mutant and WT cells were grown for 5 days in estrogen-deprived media followed by RNA extraction. Total RNA was profiled for miRNAs using the Nanostring Human v3 miRNA Assay, which examines the expression of nearly 800 miRNAs. We found that 84 miRNAs are differentially expressed in the D538G, Y537S, or both mutant cell lines compared to WT cells (Figure 1 and Supplementary Table S1).

**Figure 1.**
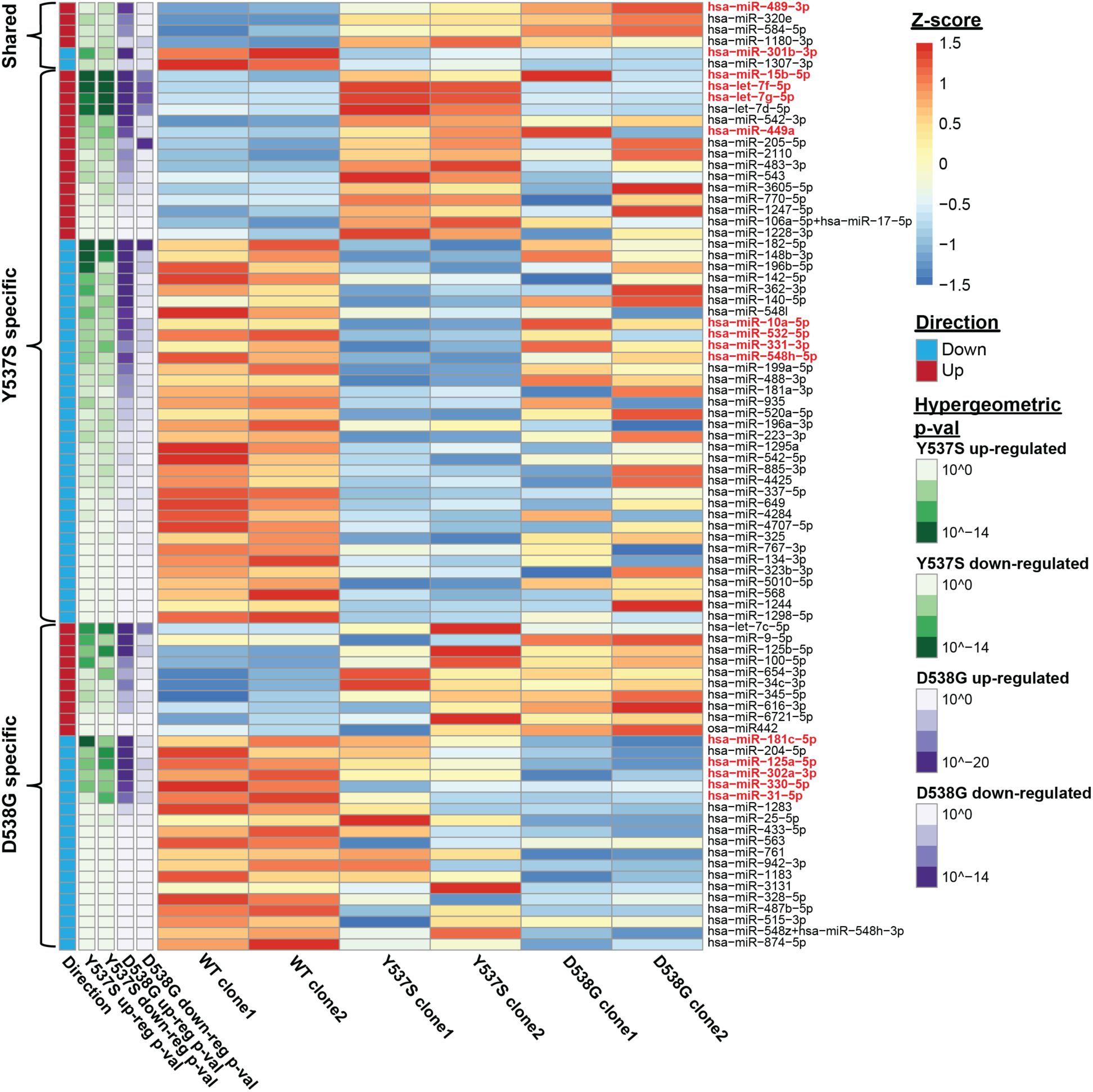
ER mutations drive the differential expression of miRNAs. Heatmap shows expression levels of 84 miRNAs identified as differentially expressed in ER mutant cells compared to WT cells. Hypergeometric p-values between each miRNA’s respective target genes and previously defined mutant-specific genes are indicated on the left. Darker color indicates more significant hypergeometric p-values. 15 miRNAs selected for further investigation are highlighted in red.

To identify which differentially expressed miRNAs likely contribute to mutant-specific gene expression, we utilized the DIANA-TarBase v8, an open-access reference database of experimentally supported miRNA targets (43). We compared the target genes for each differentially expressed miRNA with our previously identified mutant-specific genes. Because miRNAs effectively down-regulate the expression of their target genes by promoting their degradation, we compared the target genes of miRNAs that exhibited increased expression in ER mutant cells to mutant-specific down-regulated genes and the target genes of miRNAs that were down-regulated in ER mutant cells to mutant-specific up-regulated genes. Using this approach, we identified 15 differentially expressed miRNAs whose target genes are significantly enriched for mutant-specific genes (Figure 1 and Supplementary Figure S1A-E). Of these 15 miRNAs, 2 (miR-301b and miR-489) were differentially expressed in both the Y537S and the D538G mutant cells, while the remaining 13 miRNAs were differentially expressed in Y537S cells only (8 miRNAs) or in the D538G cells only (5 miRNAs). We found that genes that are both differentially expressed in ER mutant cells and are target genes of these 15 differentially expressed miRNAs are enriched for a number of cellular processes such as increased proliferation and cell migration (Supplementary Figure 1A-D). These processes were previously observed to be significantly altered in ER mutant cells. In particular, increased cell proliferation was a strong characteristic of both Y537S and D538G ER mutant cells (16).

The differential expression of these 15 miRNAs, as well as the remaining 69 differentially expressed miRNAs, occurs as a result of mutant ER expression and could be directly regulated by ER. To determine the role of ER in regulating these miRNAs we took two approaches. We first treated WT ER cells with E2 for 25 days, recapitulating long-term ER activity, and performed miRNA expression analysis. We found that only 12 of the 84 mutant ER regulated miRNAs were similarly differentially expressed upon long-term ER activation (Supplementary Figure S2A). Of the 15 miRNAs identified above, 5 were found to be regulated by long-term E2 treatment in WT cells. We next tested the role of ER in regulating miRNAs by knocking down ER expression using siRNAs. siRNA treatment reduced ER protein levels in mutant cells by over 50% (Supplementary Figure S2B). We performed miRNA expression analysis on mutant ER cells after ER knock-down and found that only 7 out of the 84 mutant-regulated miRNAs were significantly differentially regulated in the expected direction, with the low number of significant miRNAs likely the result of high clone variation (Supplementary Figure S2C). However, nearly half of the 84 miRNAs that were differentially expressed in mutant ER cells exhibited expression changes in the expected direction upon ER knock-down, suggesting that mutant ER is likely playing a direct role in the dysregulation of several miRNAs.

Using WT and mutant ER ChIP-seq data (16), we further investigated the potential contribution of mutant ER in regulating the expression of miRNAs. Of the 15 miRNAs identified above as likely contributors to mutant ER gene expression, 12 had at least one ER-binding site (ERBS) within 10kb and 13 had at least one ERBS within 100kb of their respective pri-miRNA TSS (50). Three of these miRNAs had two or more ERBS within 10kb while 12 had two or more ERBS within 100kb of their pri-miRNA TSS (Supplementary Figure S2D). Additionally, 4 of these miRNAs had ERBS within 100kb that exhibited significantly increased or decreased ER binding in ER mutant cells compared to WT cells. Of the 12 miRNAs that had at least one ERBS within 10kb of the pri-miRNA TSS, 7 were found to exhibit moderate, yet non-significant changes in expression upon ER knock-down in the expected direction. We also utilized previously obtained ATAC-seq data and found that differential chromatin accessibility near differentially expressed miRNAs may also factor into the altered expression of 6 out of 15 miRNAs (Supplementary Figure S2D). These findings suggest that the differential expression of many of these miRNAs is at least in part directly controlled by mutant ER and possibly downstream factors.

miRNAs located in the introns of protein-coding genes can be transcribed from their host gene promoter and expressed as a byproduct of the host gene transcription (51). To determine the contribution of host gene differential expression to miRNA regulation, we compared the expression patterns of differentially expressed intronic miRNAs to the expression of their host genes as observed previously (16). We found that only 4 of 30 intronic miRNAs exhibited expression patterns that coordinated with the expression of their host genes in ER mutant cells compared to WT cells (miR-489/CALCR, miR-1180/B9D1, miR-140/WWP2, and miR-15b/SMC4). These results suggest that the altered expression of most intronic miRNAs is independent of host gene regulation.

### Differentially regulated miRNAs in ER mutant cells significantly impact cell proliferation

To more directly determine the effects of differential miRNA expression on cell proliferation in ER mutant cells, we used miRNA mimics to increase a miRNA’s effect or miRNA inhibitors to decrease a miRNA’s effect in both ER WT and ER mutant cells. WT or ER mutant cells were transfected with a single miRNA mimic or inhibitor for each of the 15 differentially expressed miRNAs that exhibited enrichment for mutant-specific genes in their respective target gene sets. Immediately following inhibitor or mimic introduction, cells were monitored for proliferation for 48 hours using the IncuCyte Zoom platform for live-cell imaging. Treatment with a miRNA mimic or inhibitor led to significant changes in proliferation rates in mutant and/or WT ER cells for several of the 15 miRNAs examined, with miR-301b-3p, miR-181c-5p, and miR-331-3p producing particularly notable effects (Figure 2A-C and Supplementary Figure S3A-C).

**Figure 2.**
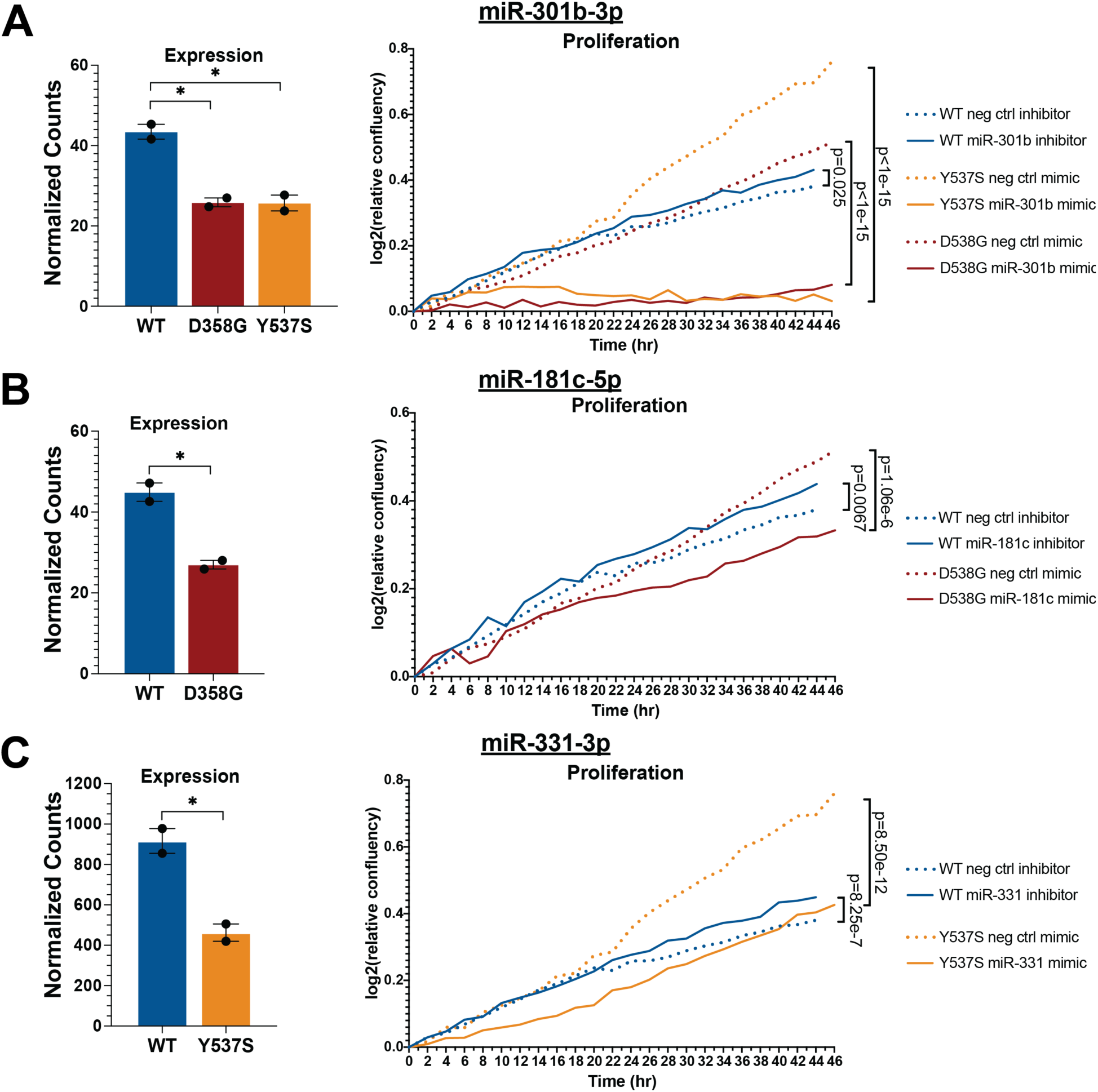
Altered miRNA activity significantly impacts proliferation in ER mutant cells. Expression levels and relative confluence are shown over time for miR-301b (**A**), miR-181c (**B**), and miR-331 (**C**). Bar graphs show normalized counts for each miRNA for ER WT, D538G, and Y537S clones. Error bars indicate average <u>±</u> SEM for two clones for each genotype. Student two-sample *t* test was used: *, P< 0.05; **, P < 0.01; ns, not significant. Proliferation graphs show log2 transformed normalized confluence values over 48 hours. Significance between the slopes of the lines was determined using a generalized linear model and the Wald test.

In Y537S and D538G ER mutant cells, where miR-301b is significantly down-regulated, treatment with a mimic for miR-301b in Y537S and D538G ER mutant cells had a striking effect, completely abrogating proliferation in both ER mutant models (Figure 2A, Supplementary Figure S3A). Treatment with a miR-301b inhibitor in mutant cells increased proliferation in Y537S ER mutant cells but had no measurable effect on D538G mutant ER cells. Conversely, treatment of WT ER cells with a miR-301b inhibitor, recapitulating the loss of miR-301b in ER mutant cells, or miR-301b mimic had negligible effects on cell proliferation. miR-301b has been shown in other contexts to drive increased proliferation (52,53). However, these results suggest that the miR-301b-mediated effects on proliferation are dependent on the cells’ ER mutation status and that loss of miR-301b contributes to increased proliferation rates specifically in ER mutant cells.

Treatment with mimics or inhibitors for the other two miRNAs of interest led to milder effects on proliferation. Introduction of a mimic of miR-181c, which was down-regulated in D538G mutant cells, significantly decreased proliferation in both D538G and WT ER cells (Figure 2B; Supplementary Figure S3A). However, further decreasing the activity of miR-181c in D538G mutant cells by treating with a miR-181c inhibitor had no significant effect on proliferation (Supplementary Figure S3A). In WT ER cells, decreasing miR-181c activity via miR-181c inhibitor treatment, recapitulating the loss of miR-181c seen in D538G cells, led to a slight yet significant increase in proliferation (Figure 2B). Increasing the activity of miR-331, which was down-regulated in the Y537S mutant, resulted in a significantly decreased proliferation rate in both Y537S mutant and WT ER cells, while treatment of Y537S mutant or WT cells with a miR-331 inhibitor increased proliferation in both cell lines, although the effect in WT cells was moderate (Figure 2C; Supplementary Figure S3A). The mimic or inhibitor treatments for the remaining 11 miRNAs, which were organized into 5 groups based on their expression patterns in WT and ER mutant cells, resulted overall in weaker proliferative effects (Supplementary Figure S3B-C). Together, these data show that multiple miRNAs differentially expressed in ER mutant cells contribute to the regulation of cell proliferation, a phenotype that is increased in ER mutant cells. In the ER mutant setting, these miRNAs changes drive increased proliferation with some miRNAs having a stronger effect on proliferation than others.

### Differentially expressed miRNAs contribute to mutant-specific gene regulation

We investigated the effects of miR-301b, miR-181c, and miR-331, which impacted proliferation, as well as let-7f, which was exceptionally up-regulated in Y537S mutant cells, on gene expression in ER mutant cells by performing RNA-seq on total RNA collected from ER mutant or WT cells after treatment with a single miRNA mimic or inhibitor. miRNAs that were down-regulated were evaluated with a miRNA mimic (miR-301b-3p, miR-181c-5p, miR-331-3p), while let-7f was evaluated with a miRNA inhibitor. In this way, the activity of the each miRNA of interest was effectively reverted towards WT levels. RNA-seq was performed 24 hours post-transfection.

We found that altered expression of these miRNAs by mimic or inhibitor treatment resulted in the differential expression of tens to hundreds of genes (miRNA-altered genes) depending on the miRNA (Figure 3A and Supplementary Table S2). To validate that the observed gene expression changes are a result of mimic or inhibitor treatment, we asked whether miRNA-altered genes exhibited enrichment for target genes of the corresponding miRNA. Mimic treatment should lead to enrichment of miRNA targets in down-regulated genes, while inhibitor treatment should exhibit enrichment of miRNA targets in up-regulated genes. As expected, we observed an enrichment for predicted miRNA target genes in down-regulated genes altered upon miR-301b, miR-181c, or miR-331 mimic treatments and in up-regulated genes after let-7f inhibitor treatment (Supplementary Figure S4A). We found that inhibition of let-7f resulted in very few gene expression changes (n = 16), potentially due to the high expression of this miRNA in Y537S ER cells, although the affected genes were nearly all previously identified as let-7f target genes. The enrichment for miR-331 targets in down-regulated genes after miR-331 mimic treatment was relatively weak. However, more miR-331 target genes were observed in the group of down-regulated genes than in the group of up-regulated genes suggesting that the differential gene expression observed after mimic treatment may still be driven by altered miR-331 activity and that the target genes of miR-331 may not be well defined or could be highly cell type specific.

**Figure 3.**
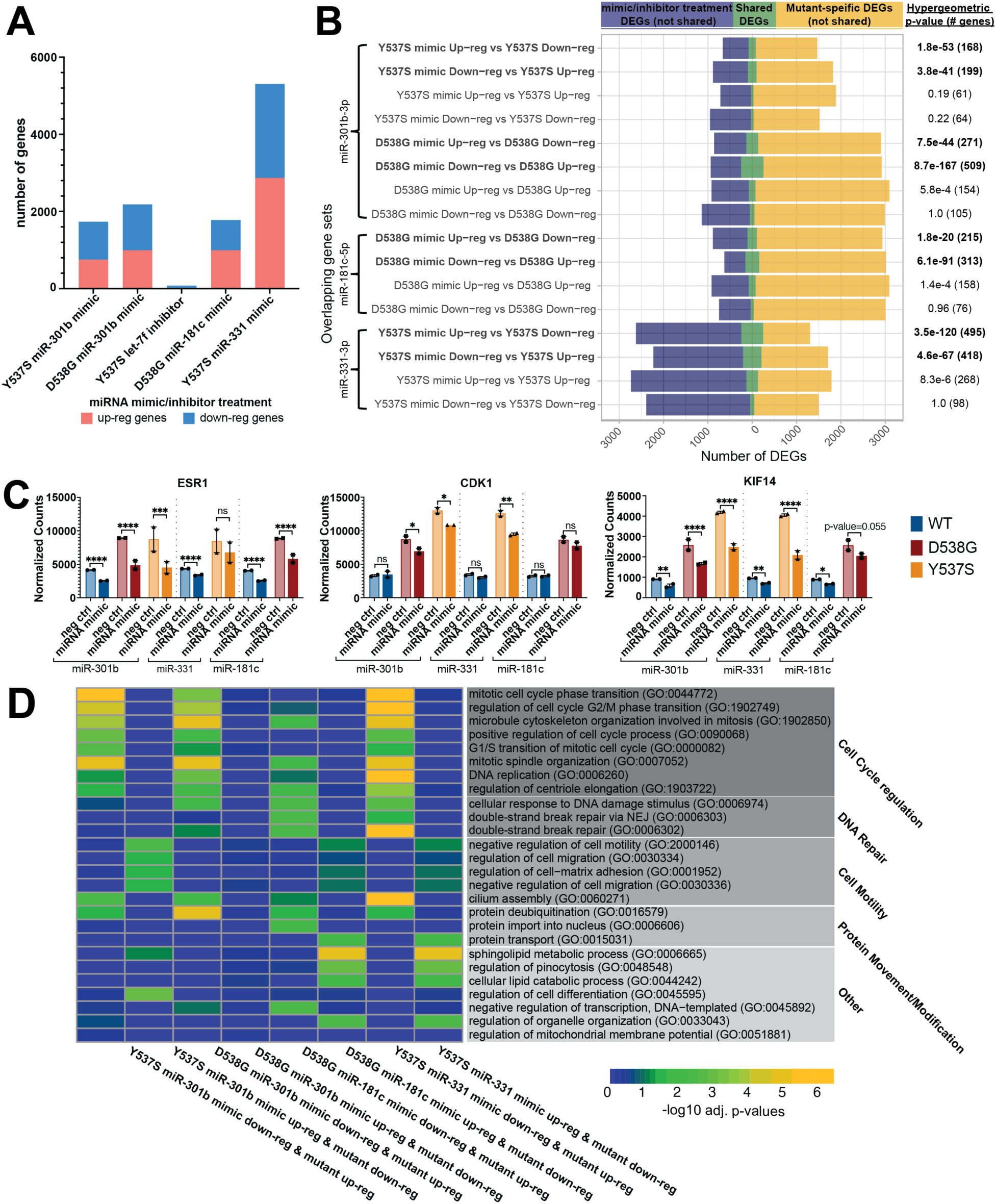
Altered miRNA activity leads to differential expression of many mutant-specific genes involved with processes driving cancer-associated phenotypes. (A) Total number of genes altered upon miRNA mimic or inhibitor treatment is shown; red indicates up-regulated genes and blue indicates down-regulated genes. (B) Overlap (green) of miRNA mimic-or inhibitor-treated differentially expressed genes (DEG) (blue) and mutant-specific DEG (yellow) is displayed for each gene set overlap. Significance of overlapping gene sets, indicated on the right, are hypergeometric p-values. (C) Examples of miRNA-altered mutant-specific genes are shown. Error bars represent average <u>±</u> SEM for two clones for each genotype. Adjusted p-values are indicated as calculated by DESeq2 (Wald test): *, P< 0.05; **, P < 0.01; ***, P < 0.001; ****, P < 0.0001; ns, not significant. (D) Heatmap indicates the enrichment for miRNA-altered mutant-specific genes in the indicated gene ontology terms. Colors indicate the -log10 adjusted p-values.

To determine whether our miRNAs of interest contribute to the mutant-specific gene expression patterns we observed previously (16), we asked whether miRNA-altered genes were significantly enriched for mutant-specific genes. Because mimic or inhibitor treatment reversed the activity of these miRNAs relative to their expression in ER mutant cells compared to WT cells, we expected to find significant overlaps between mimic-or inhibitor-regulated genes and mutant-specific genes regulated in the opposite direction. For example, we would expect genes down-regulated by miR-301b mimic treatment to be up-regulated in ER mutant compared to WT cells. As expected, we found that genes that were up-regulated upon mimic or inhibitor treatment were significantly over-represented in the corresponding mutant-specific down-regulated gene sets (Figure 3B). Conversely, genes that were down-regulated upon mimic or inhibitor treatment were significantly enriched for mutant-specific up-regulated genes. Comparison of mutant-specific genes regulated in the same direction as miRNA-altered genes (i.e. mutant-specific up-regulated vs miRNA mimic/inhibitor up-regulated) resulted in non-significant or very low enrichment between gene sets (Figure 3B). These results show that miRNAs regulate the expression of many mutant-specific genes in the expected direction and suggest that they impact gene regulation and proliferation in ER mutant cells.

The majority of mutant-specific gene expression changes can be explained by mutant ER’s constitutive activity, altered ER genomic binding, or altered chromatin accessibility (16). However, the mechanism behind the differential expression of the remaining mutant-specific genes was unexplained. Our findings showed that miRNA-altered mutant-specific genes are represented in each of these categories at a distribution similar to that observed for all mutant-specific genes (Supplementary Figure S4B). miRNAs contribute considerably to these previously-unexplained mutant-specific gene expression effects. We found that 4 miRNAs (miR-301b, miR-181c, miR-331, and let-7f) together could explain between 18% and 34% of mutant-specific genes that were unexplained from our previous findings (D538G: 18.7%; Y537S: 33.2%; sup. table S3), indicating that miRNAs could be playing a substantial role in shaping ER mutant-driven gene expression patterns.

We found that one of the mutant-specific genes regulated by miRNAs is *ESR1*, the gene encoding ER. miR-301b and miR-181c were both found to regulate ER, which has been shown to have complementary sequence in its 3 UTR to the seed sequences for both of these miRNAs (Figure 3C; Supplementary Figure S4C) (48,54). ER is significantly up-regulated in ER mutant cells, where ER is constitutively active, compared to WT cells (Figure 3C). This is counter to some reports that show that ER, in the mammary gland, regulates its own expression through a negative feedback mechanism in which ER activity leads to decreased transcript levels of *ESR1* (55–57). This suggests that ER mutant cells are able to evade the negative regulation typically exerted by active ER and potentially take advantage of the resulting sustained or even heightened levels of ER to promote increased cell growth and proliferation. The observation that multiple miRNAs targeting *ESR1* are significantly down-regulated in ER mutant cells indicates an ER mutant-driven down-regulation of miRNAs as a key mechanism for evading normal negative feedback regulation by active ER on *ESR1* expression. Consistent with our observations, we previously found that metastatic breast cancer patient samples harboring D538G or Y537S mutant ER exhibit significantly increased levels of ER compared to WT ER breast cancer metastases (17).

To examine the overall effects of miRNAs in regulating ER mutant phenotypes, we performed gene ontology analysis on miRNA-altered mutant-specific genes. This analysis revealed that these genes play roles in several important biological processes that are altered in mutant cells. miR-301b and miR-331 altered mutant-specific up-regulated genes were enriched for processes related to cell cycle progression (Figure 3D). Many of these genes, such as *CDK1* and *KIF14*, play important roles in mediating cell cycle progression, indicating again the role of these miRNAs in driving increased proliferation (Figure 3C). miRNA-altered mutant-specific up-regulated genes resulting from miR-181c mimic treatment in D538G cells were also involved in cell cycle regulation, although these genes were more enriched in DNA damage repair (Figure 3D). Additional gene ontology terms associated with miRNA-altered mutant-specific genes include cell-matrix adhesion and negative regulation of cell motility (Figure 3D). These genes were down-regulated in ER mutant cells and up-regulated in cells treated with either a miR-301b, miR-181c, or miR-331 mimics, suggesting that these miRNAs restrict cell motility. The decreased expression of these miRNAs in ER mutant cell lines may contribute to increased cell motility and migration observed in ER mutant cells (9,14,17,58,59).

To understand the direct effects of miRNAs in driving these broad phenotypic effects, we asked which miRNA-altered mutant-specific genes are predicted direct miRNA targets based on the miRDB database for miRNA target genes, which uses an algorithm trained on large RNA-seq and CLIP-ligation data sets to predict miRNA targets (48). We found that miRNA-altered mutant specific genes were enriched for miRNA target genes in the expected direction. Of the genes overlapping mutant-specific up-regulated genes, 23% of miR-301b mimic (p-value = 1.7e-26 (Y537S) and 1.2e-64 (D538G); hypergeometric test) and 48% of miR-181c mimic (p-value = 2.1e-108; hypergeometric test) down-regulated genes were direct miR-301b or miR-181c target genes (Figure 4A, Supplementary Figure S5A). All of the let-7f inhibitor (p-value = 1.0; hypergeometric test) up-regulated genes shared with mutant-specific down-regulated genes were let-7f direct targets (Supplementary Figure S5A). However, only 1.5% (pvalue = 0.45; hypergeometric test) of miR-331 mimic down-regulated mutant-specific up-regulated genes were direct miR-331 target genes, similar to the lack of miR-331 target gene enrichment described above (Supplementary Figure S5A). We found that direct target genes of these miRNAs were enriched for several key cell signaling pathways including PI3K signaling, nuclear receptor signaling, and EGF receptor signaling, specifically through the MAPK signaling cascade (Figure 4B; Supplementary Figure S5B).

**Figure 4.**
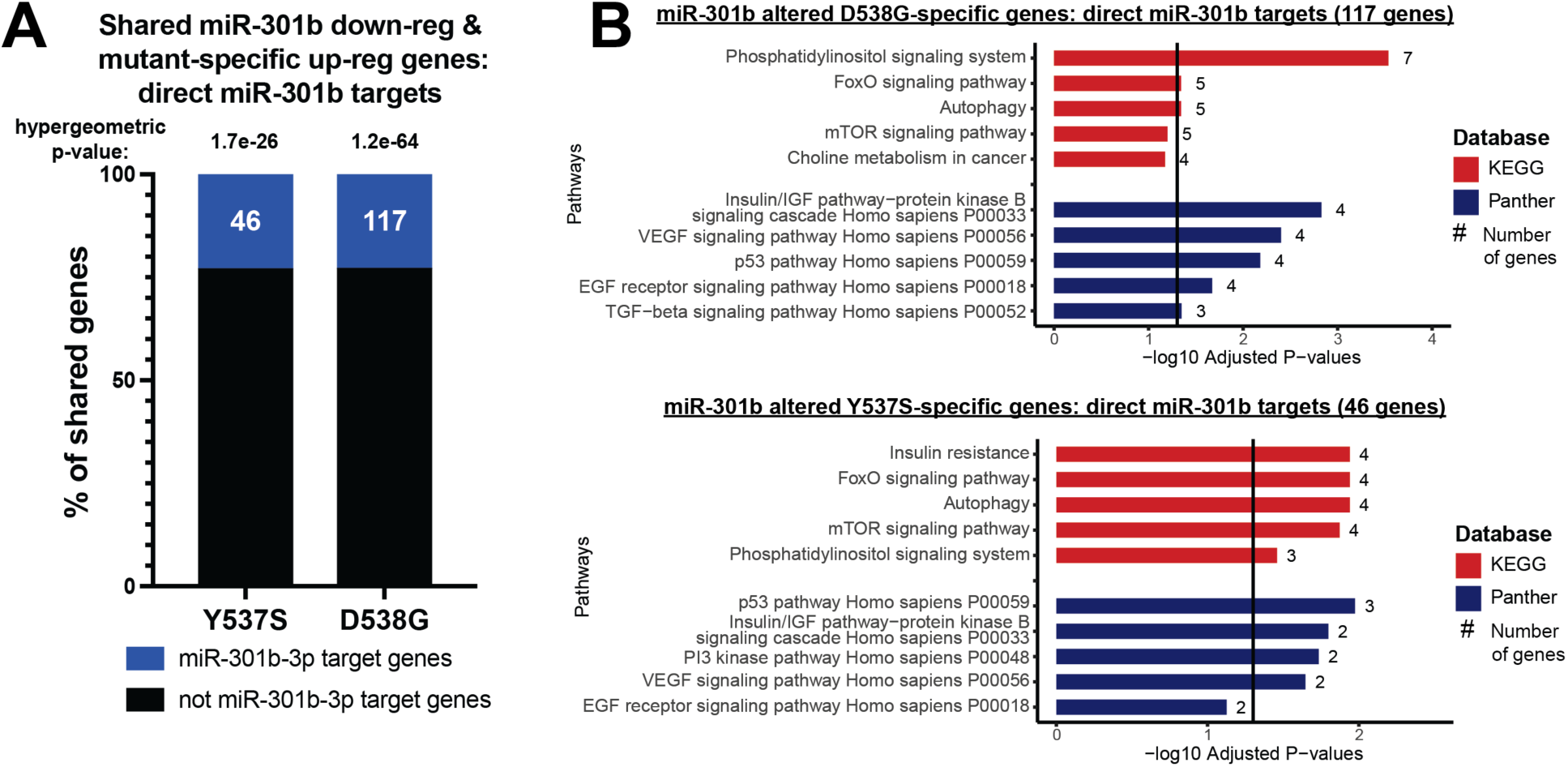
miRNAs in ER mutant cells directly target mutant-specific genes involved in important cell signaling pathways. (**A**) Percent of miR-301b mimic down-regulated mutant-specific up-regulated genes that are predicted direct targets of miR-301b is shown. Hypergeometric tests were used to calculate p-values. (**B**) Bar graphs show pathways found to be enriched in miRNA-altered mutant-specific genes. -log10 adjusted p-values are indicated and the number of genes in each pathway gene set are shown.

### miR-301b target PRKD3 contributes to increased proliferation in ER mutant cells

Analysis of miR-301b direct target genes that were also mutant-specific genes found that many of these shared genes are key components of major cell signaling pathways including MAPK signaling, PI3K signaling, and ER signaling. Of note, miR-301b targets *ERK2*, a central component of MAPK signaling which is known to drive cell proliferation, differentiation, and survival (Figure 5A), reviewed in (60). miR-301b also targets PIK3CB (P110-β), the catalytic subunit of PI3Kβ, which has been shown to enhance cell proliferation and promote tumorigenesis in breast cancer (Figure 5A), reviewed in (61). The regulation of these pathway components by differentially expressed miRNAs in ER mutant cells indicate that altered miRNA expression in ER mutant cells directly alters important cellular pathways with roles in regulating cell growth and proliferation.

**Figure 5.**
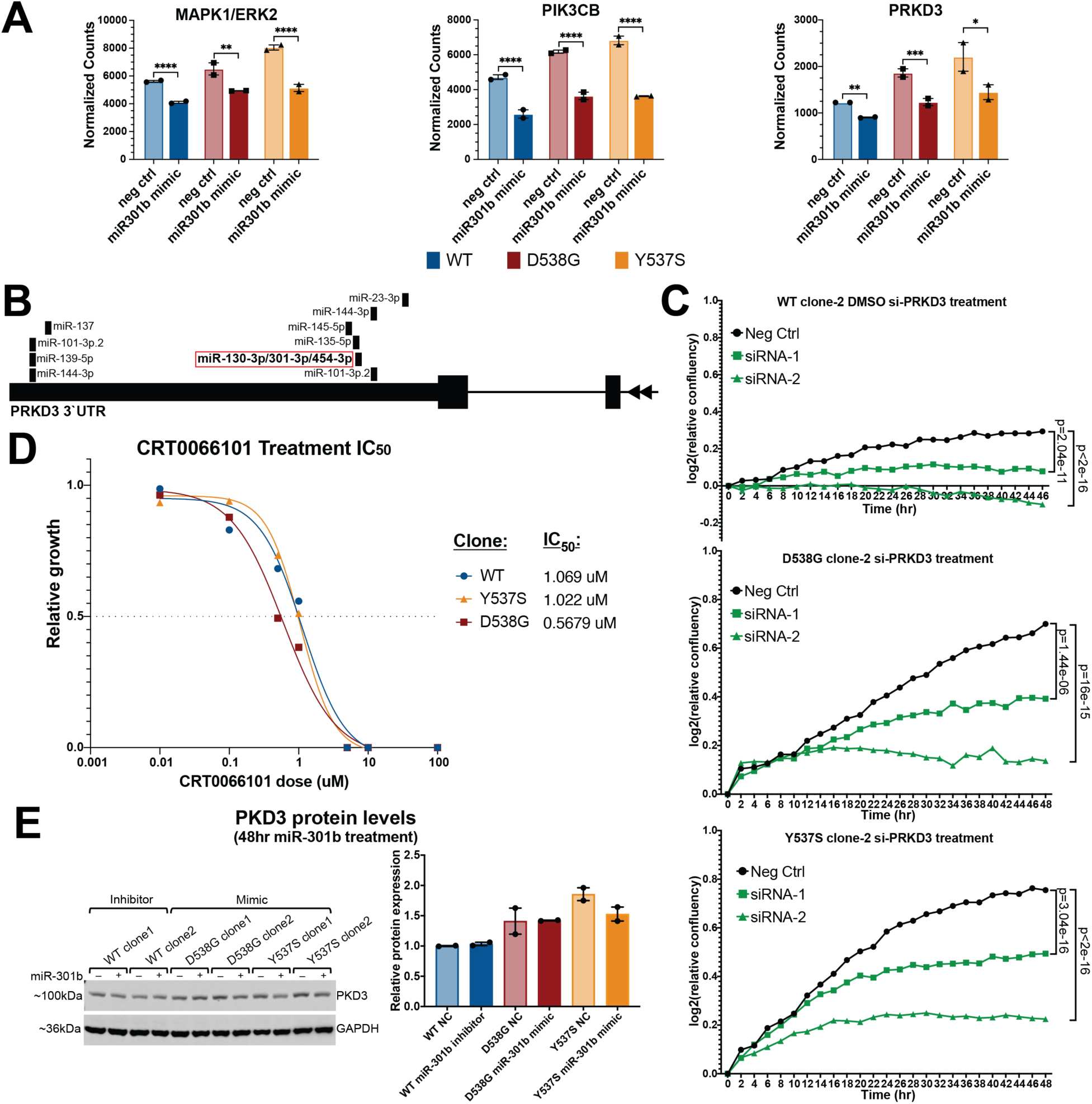
Inhibition of *PRKD3* (PKD3) reduces proliferation in ER WT and mutant cells. (**A**) Bar graphs show three examples of miR-301b altered mutant-specific genes that are likely direct miR-301b target genes, including *PRKD3*. Error bars represent average ± SEM for two clones for each genotype. Adjusted p-values are indicated as calculated by DESeq2 (Wald test). (**B**) Diagram of *PRKD3* 3’UTR indicates the regions with miRNA target sequences. The miR-301b target site is shown in the red box. (**C**) Line graphs show log2 transformed relative confluence over time of WT or mutant ER cells treated with anti-*PRKD3* siRNAs or negative control. Significance between the slopes of the lines was determined using a generalized linear model and the Wald test. (**D**) Relative growth is shown for WT, Y537S, and D538G ER clones after treatment with CRT0066101 (pan-PKD inhibitor). (**E**) Immunoblot analysis shows PKD3 expression after miR-301b inhibitor treatment in ER WT cells and miR-301b mimic in ER mutant cells. Bar graph shows quantification of PKD3 signal for two clones for each ER genotype. Error bars indicate average signal ± SEM.

miR-301b also regulates the RNA levels of *PRKD3* (the gene encoding protein kinase D3 – PKD3), which regulates activity of several signaling pathways by phosphorylating key pathway components leading to increased proliferation and cell migration (39–42). *PRKD3* gene expression is significantly up-regulated in both Y537S and D538G ER mutant cells but is significantly down-regulated upon treatment with a miR-301b mimic (Figure 5A). *PRKD3* contains a recognition sequence for miR-301b in its 3’UTR indicating that it is likely a miR-301b target gene (Figure 5B) (54). Based on these observations, we assessed whether *PRKD3* is important for proliferation of ER mutant cells in a similar manner to miR-301b mimic treatment. We knocked down *PRKD3* expression, using siRNAs, and found that treatment with either of two anti-*PRKD3* siRNAs significantly reduced *PRKD3* expression and cell proliferation in ER mutant and WT cells (Figure 5C; Supplementary Figure S6A-B). Because knock-down of *PRKD3* via siRNA reduced proliferation, we investigated whether pharmacological inhibition of PKD3 with CRT0066101 could similarly reduce cell proliferation. CRT0066101 is a pan-PKD inhibitor that has been shown to reduce proliferation in a number of tumor types, including ER-negative breast cancer, pancreatic cancer, and colon cancer (62–65). We treated ER mutant and WT cells with CRT0066101 at a range of doses and found that CRT0066101 treatment reduced cell proliferation in all cell lines in a dose-dependent fashion (Figure 5D; Supplementary Figure S6C). Significant reduction in proliferation rates was seen at concentrations as low as 0.1 uM, with IC50 values between 0.5 uM and 1.1 uM depending on ER mutation status, similar to IC50 values observed in other successful treatment settings (62,64). Significant loss of proliferation upon knock-down or inhibition of *PRKD3* points to *PRKD3* as a promising therapeutic target in ER+ breast cancers, including those containing ER mutations.

Unexpectedly, we found that while miR-301b perturbations impacted RNA levels of *PRKD3*, we only observed minimal effects on PKD3 protein levels with miR-301b mimics or inhibitors (Figure 5E). It is unclear why the effects on *PRKD3* RNA levels do not correspond to PDK3 protein levels, although this finding suggests that PKD3 protein may not play an important role in the miR-301b growth phenotype. However, the results are still consistent with PKD3 representing an important factor that promotes ER mutant and WT breast cancer cell growth.

### Differential expression of miRNAs in mutant ER cell lines is recapitulated in patient-derived xenograft models

To determine if the differential expression of miRNAs seen in the cell line models is observed in additional models, we used patient-derived xenograft (PDX) breast cancer models (66), harboring either WT ER or naturally-occurring heterozygous Y537S or L536P ER mutations. PDX models included two WT ER lines, a series of three L536P ER mutant lines from the same patient, and two Y537S ER mutant lines from the same patient. Results of miRNA profiling revealed that over 400 miRNAs were significantly differentially expressed between the Y537S and WT PDX lines, while nearly 60 miRNAs were significantly differentially expressed in the L536P PDX lines vs WT (Figure 6A; Supplementary Table S1). When compared to miRNAs that are differentially expressed in the T-47D ER mutant cell lines, only one up-regulated miRNA was shared between the Y537S PDX lines and the T-47D Y537S cell lines. However, half of the down-regulated miRNAs identified in T-47D Y537S mutant ER cell lines were also significantly down-regulated in Y537S PDX lines (Figure 6B). Of the 15 miRNAs that were differentially expressed in T-47D cells and exhibited significant overlap of target genes with mutant-specific genes, 10 were differentially expressed in the T-47D Y537S mutant clones. Of these, 5 exhibited significant up-or down-regulation in the Y537S PDX lines in the same direction as that observed in the T-47D Y537S mutant clones, including miR-301b and miR-331, while an additional 3, including miR-181c and let-7f, were trending in the same direction but were below the significance threshold (Figure 6C-D). miRNA let-7f-5p was up-regulated with a nearly significant p-value, while miR-181c was not differentially regulated in T-47D Y537S mutant cell lines and therefore was not expected to be differentially regulated in Y537S PDX lines. Of the 5 miRNAs that were differentially regulated in the T-47D D538G mutant cell lines only, none were differentially regulated in the PDX Y537S lines, indicating consistent ER allele-specific effects. These miRNA expression results in WT and mutant ER PDX lines show that miRNAs are significantly altered in patient-derived models harboring mutant ER and validate many of our findings in T-47D cells.

**Figure 6.**
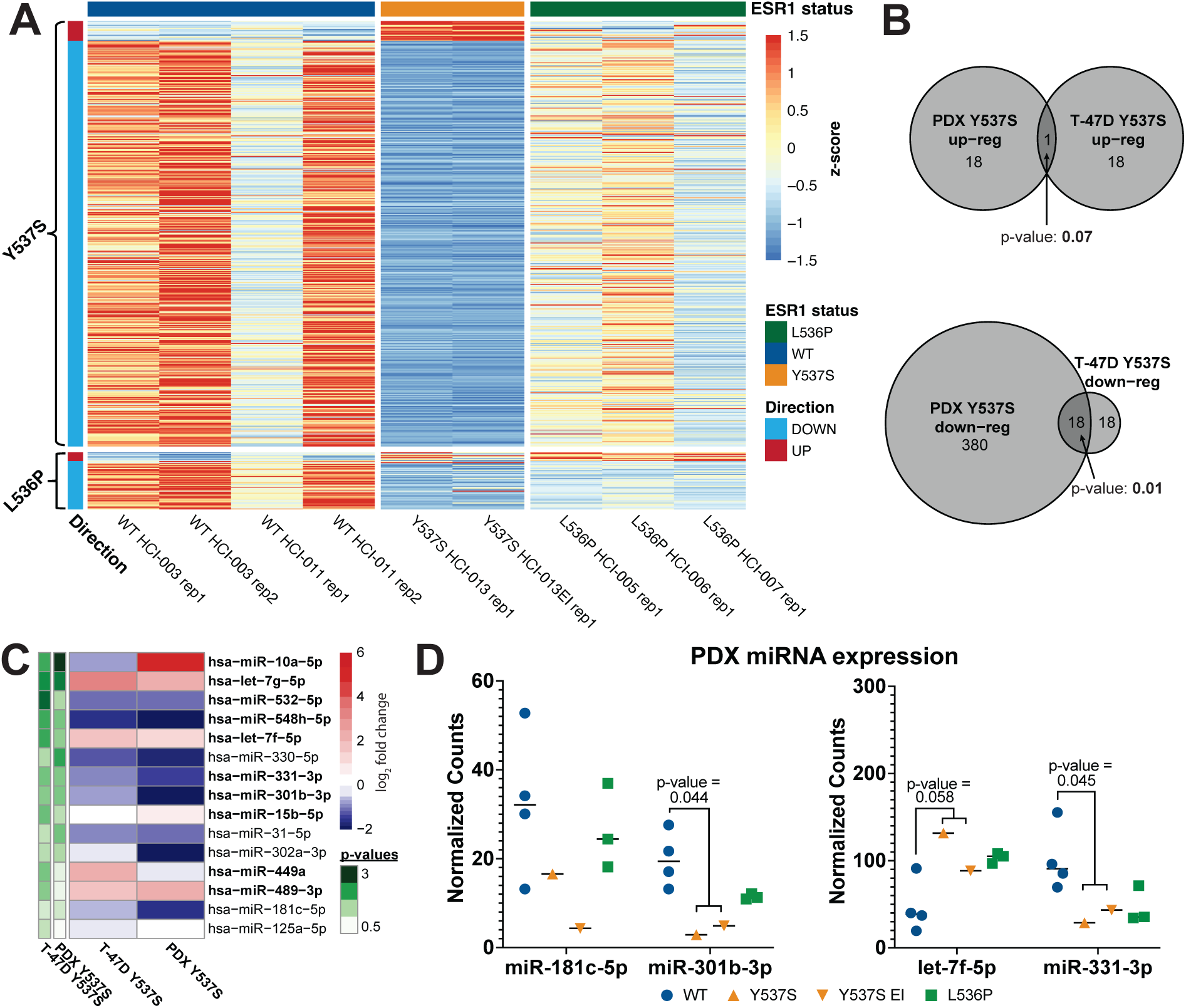
miRNAs are altered in PDX models similar to changes observed in cell line models. (**A**) Heatmap representing differentially expressed miRNAs in L536P or Y537S ER mutant PDX models compared to WT ER models. (**B**) Overlap between PDX and T-47D differentially expressed miRNAs. Significance was determined using a hypergeometric test. (**C**) Heatmap of the 15 differentially expressed miRNAs analyzed in T-47D clones for proliferation effects. Heatmap shows the log2 fold change values for these miRNAs in T-47D and PDX Y537S models. Bolded miRNAs were differentially expressed in T-47D Y537S models. -log2 p-values are indicated in green. (**D**) Expression of miR-181c, miR-301b, let-7f, and miR-331 in L536P and Y537S PDX models. P-values were calculated using a Student’s two sample *t* test.

## Discussion

The frequent occurrence of ER mutations in hormone therapy-resistant ER-positive metastatic breast cancer has led to intense research efforts to elucidate the molecular and phenotypic effects of ER mutations and to identify vulnerabilities in ER mutant cancers that could be exploited to develop effective therapies for hormone therapy-resistant tumors (4–18). In a previous study, we used isogenic T-47D models expressing the Y537S or D538G ER mutations from the endogenous ER locus to show that ER mutations regulated the differential expression of thousands of genes and that these transcriptional changes were accompanied by phenotypic changes, including increased migration, invasion, and proliferation (16,17). Although our understanding of ER mutations and their effects has improved significantly, the contribution of miRNAs to ER mutant gene regulation and downstream effects was unknown. In humans, over 2000 miRNAs regulate the expression of roughly 60% of protein coding genes (67,68), indicating a potential for miRNAs to aid mutant ER in alterating gene expression. In this study, we aimed to determine the role of miRNAs in regulating transcriptional and phenotypic changes in mutant ER breast cancer.

Using the isogenic cell line models introduced previously (16), we found that 84 miRNA (over 10% of those profiled) were differentially expressed in the Y537S, D538G, or both mutant cell lines. Mutant-specific genes identified in our previous work were over-represented in the target genes of many of these miRNAs. In several cases, namely miR-301b, miR-181c, miR-331, and let-7f, these miRNA targets were enriched for genes with important roles in driving cell growth, proliferation, or motility, indicating a possible role for these miRNAs in driving cancer-associated phenotypes that we observed previously in ER mutant cells. When we directly tested the effect of these miRNA on ER mutant-specific gene expression by altering miRNA activity through mimic or inhibitor treatment followed by RNA-seq, we found that these miRNAs did indeed alter the expression of many mutant-specific genes. Furthermore, these miRNA-altered mutant-specific genes were enriched for cell cycle-related genes and therefore altered activity of these miRNAs in ER mutant cells can partially explain the observed increased proliferation.

Altered expression of miRNAs in ER mutant cells can lead to broad gene expression changes both through direct miRNA targeting or by downstream effects in which miRNAs directly alter transcriptional regulators which in turn alter the expression of their target genes. We found that for three of the four miRNAs we investigated in depth (miR-301b, miR-181c, and let-7f), mutant-specific genes that were up-or down-regulated upon miRNA inhibitor or mimic treatment, respectively, were highly enriched for predicted direct target genes of the respective miRNA. For miR-301b and miR-181c, many direct target genes were components or regulators of important cell signaling pathways. Altered expression of these pathways could lead to additional gene expression changes, potentially explaining the remaining 50-75% of miRNA-altered mutant-specific genes. Only 2 of the 17 genes up-regulated by let-7f inhibitor treatment were also mutant-specific down-regulated genes, but both of these genes (TUSC2 and PLAGL2) were predicted let-7f direct target genes. These findings indicate that miRNA-mediated mutant-specific gene expression changes occur as a result of both direct miRNA-mediated mRNA degradation and through secondary effects driven by altered expression or activity of genes that are direct miRNA targets.

One major question regarding differentially expressed miRNAs is what drives their altered expression. Altered miRNA expression could be driven directly by ER, by other transcription factors downstream of ER signaling, or by other transcription factors that have improved or decreased access to genomic regions due to mutant ER driven changes in chromatin accessibility (16). miRNA expression analysis following long-term E2 treatment in WT ER cell lines indicated that less than 15% of mutant ER regulated miRNAs are regulated by long-term activation of WT ER. Furthermore, ER knock-down by siRNAs showed that while only a small percentage (8.3%) of mutant ER regulated miRNAs are significantly differentially expressed upon ER loss via siRNA treatment, nearly half of mutant ER regulated miRNAs change expression in the expected direction after ER knock-down (D538G: 42.9%; Y537S: 49.1%). Utilizing mutant ER ChIP-seq data and ATAC-seq data from our previous study, we found altered ER binding near the pri-miRNA TSSs for let-7f, miR-301b, miR-331, and miR-181c, indicating direct regulation of these miRNAs by mutant ER. Overall, our results indicate that mutant ER can impact miRNA expression both directly and indirectly.

Not only does mutant ER likely regulate the expression of these miRNAs, but conversely, at least two differentially expressed miRNAs (miR-301b and miR-181c) regulate the expression of *ESR1*, the gene encoding ER. miR-301b and miR-181c are down-regulated in ER mutant cells which correlates with increased expression of ER. We found that reversing the low expression of these miRNAs to higher levels, such as those observed in WT cells, through miRNA mimic treatment, resulted in significantly decreased *ESR1* expression. In addition, *ESR1* contains recognition sequences for both miR-301b and miR-181c in its 3’UTR, indicating that *ESR1* is a direct target of these miRNAs. In the normal mammary gland, ER activity results in a negative feedback loop that reduces ER expression (56). In T-47D cells, initial ER activity increases ER expression, but with long-term ER activity, ER expression plateaus and drops back towards starting levels. Mutant ER-driven down-regulation of miR-301b and miR-181c represents a mechanism by which mutant ER evades this normal negative feedback on ER expression and instead achieves higher ER levels.

In addition to understanding the role of miRNAs in regulating mutant-specific gene expression, we also aimed to understand how miRNAs contribute to cancer-associated phenotypes in ER mutant cells. One major phenotype observed in ER mutant cells is their increased proliferative capacity (16) and miRNAs could contribute to the positive effect mutant ER has on proliferation. When we tested the effect of 15 differentially expressed miRNAs on proliferation by individually modulating their activity with miRNA mimics or inhibitors, we found that several miRNAs significantly altered proliferation. In particular, miR-301b, miR-181c, and miR-331 significantly altered ER mutant cell growth. This effect is explained by the finding that the mutant-specific genes that are altered by these miRNAs include many cell cycle-related genes and genes involved in pathways that promote cell proliferation. Thus, altered expression of miRNAs driven by ER mutations likely represents an important mechanism by which ER mutant cells achieve increased proliferation.

Another possible explanation for the increased proliferation in ER mutant cells is that altered miRNA expression impacts the activity of important signaling pathways that promote proliferation. miR-301b, which is down-regulated in ER mutant cells, directly targets the *PRKD3* gene, which codes for protein kinase D3 (PKD3), a regulator of cell signaling pathways that promote cell proliferation. *PRKD3* is up-regulated in a mutant-specific fashion and altering the activity of miR-301b significantly altered the RNA levels of *PRKD3*. PKD3 promotes phosphorylation of several components of cell signaling pathways which drive proliferation in breast and other cancer types (39–42). We found that increased proliferation in ER mutant cells can be significantly reduced when PKD3 is inhibited either genetically by siRNA knock-down, or pharmacologically by treatment with the PKD inhibitor CRT0066101. We were surprised to find that miR-301b perturbations did not significantly impact protein levels of PKD3. The lack of correlation between RNA and protein is commonly observed in the human proteome and could be due to several possibilities (69). For example, reduction in *PRKD3* RNA by miR-301b might be compensated by an increase in translation efficiency or an increase in PDK3 half life. Regardless of the mechanism of protein compensation, the lack of miR-301b effects on PKD3 protein levels indicates that *PRKD3* targeting is unlikely the driving force behind miR-301b’s anti-proliferative ability. Even though miR-301b may not be primarily working through PKD3, our findings show that PKD3 inhibition is a promising option for future therapeutics, with CRT0066101 having already been shown to inhibit tumor growth both *in vitro* and *in vivo* in several other cancer types (62–65).

miRNAs that are differentially expressed in ER mutant cells could produce varying effects that are dependent on ER mutant status. miR-301b, for example, has been shown in other settings, including triple negative breast cancer, to promote proliferation (52,53). However, we found that it has the opposite effect in ER mutant cells where increasing its activity completely abolished proliferation. These findings not only provide evidence of a significant role for miRNAs in establishing ER mutant effects, but also indicate that some miRNAs are utilized in unique fashion in ER mutant cells. The effect of miRNAs is highly cell context-dependent and this observation could be explained by the altered transcriptional profile seen in ER mutant cells compared to WT cells. The differential effects could result in a different set of miR-301b target genes expressed in ER mutant cells than are expressed in WT cells, leading to a different overall effect of miR-301b in these cells.

We validated the mutant ER-driven differential expression of miRNAs by identifying differentially expressed miRNAs in PDX models of mutant or WT ER-positive breast cancer. We found that many of the same, as well as many additional, miRNAs are differentially expressed in Y537S PDX models as those identified in our T-47D cell line models. Of note, miR-301b, miR-331, and let-7f were all differentially expressed in the same direction in PDX Y537S models as in T-47D cells. This consistency between cell line and PDX models demonstrates that differential expression of miRNAs has the potential to play a significant role in mutant ER pathology in patients.

Overall our findings show that certain miRNAs exhibit consistent differential expression in mutant ER breast cancer models and are capable of regulating key genes and pathways that can be targeted to successfully inhibit breast cancer growth. These results also help to identifying additional factors and signaling pathways that could be effective drug targets to treat hormone therapy-resistant ER mutant breast cancers. Additionally, the use of miRNA mimics and inhibitors to directly alter the activity of a specific miRNA has recently become an exciting new therapeutic option currently being explored. Our finding that many miRNAs are differentially regulated in the ER mutant breast cancer setting suggests that the direct targeting of these miRNAs through mimics and inhibitors could also represent a promising therapeutic approach in treating this hormone therepy-resistant disease.

## Data Availability

All raw and processed sequencing data generated in this study have been submitted to the NCBI Gene Expression Omnibus (GEO; http://www.ncbi.nlm.nih.gov/geo/) under the following accession number: GSE214580.

## Acknowledgements

This work was supported by a DOD Breast Cancer Research Program Expansion Award, BC181341, to JG. Research reported in this publication utilized the High-Throughput Genomics Shared Resource at Huntsman Cancer Institute at the University of Utah and was supported by the National Cancer Institute of the National Institutes of Health under Award Number P30CA042014. We thank members of the Gertz, Welm, and O’Connell labs for their helpful comments on the study and the manuscript.

## Author Contributions

S.A., Z.B., and J.G. conceived the study and designed the experiments. S.A., J.T.P, K.S.O., and J.G. performed experiments, analyzed data, interpreted data, and wrote the manuscript. S.A., J.T.P, Z.B., K.S.O., A.L.W., R.M.O., and J.G. reviewed and edited the manuscript. J.G., A.L.W, and R.M.O. supervised the study.

