## Supplemental Figures and Tables S3 and S4 for "Estrogen receptor alpha mutations regulate gene expression and cell growth in breast cancer through microRNAs"

### SUPPLEMENTARY FIGURES

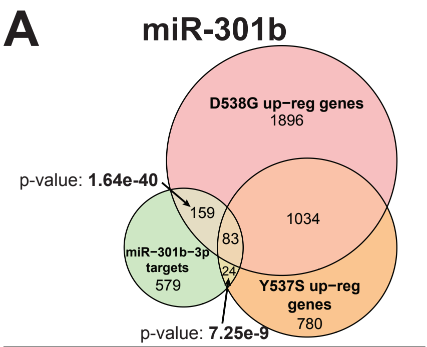


| Pathway/GO term | Y537S Adj. p-value | D538G Adj. p-value |
|---|---|---|
| Cell Cycle | 1.640e-8 | 3.830e-3 |
| G2-M Checkpoint | 1.994e-6 | 2.169e-5 |
| E2F Targets | 1.994e-6 | 4.734e-4 |
| Mitotic Spindle | NS | 2.169e-5 |
| Retinoblastoma gene in cancer | 3.859e-6 | 1.189e-2 |
| FOXM1 transcription factor network | 1.529e-5 | 2.760e-2 |
| Regulation of Microtubule Cytoskeleton | 2.677e-2 | 4.901e-3 |
| Rac 1 cell motility signaling pathway | NS | 5.663e-3 |
| Myc Targets V1 | 4.882e-2 | 8.006e-3 |

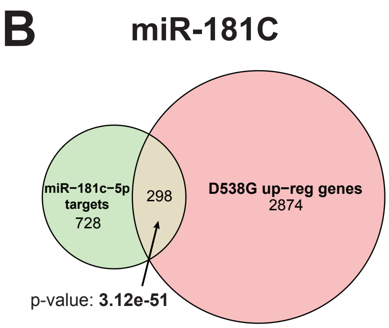


| Pathway/Go term | Adj. p-value |
|---|---|
| Gene Expression | 9.152e-5 |
| ErbB1 downstream signaling | 7.213e-5 |
| Integrated breast cancer pathway | 1.327e-4 |
| E2F Targets | 4.344e-4 |
| Mitotic Spindle | 6.971e-4 |
| Epithelial Mesenchymal Transition | 6.971e-4 |

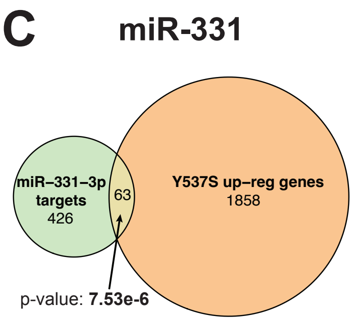


| Pathway/Go term | Adj. p-value |
|---|---|
| Mitotic Prometaphase | 1.080E-6 |
| Cell Cycle | 1.678E-6 |
| RHO GTPases Activate Formins | 2.975e-5 |
| Organelle biogenesis and maintenance | 7.251e-5 |
| Signaling events mediated by HDAC class II | 2.323e-4 |
| Integrin-linked kinase signaling | 3.846E-4 |
| Protein Nuclear Import and Export | 1.288E-3 |
| Focal adhesion | 3.583E-3 |

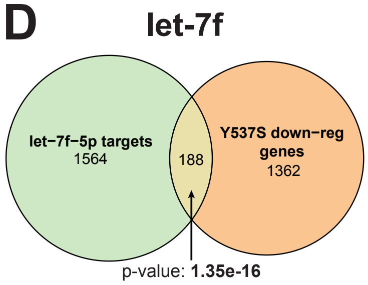


| Pathway/GO terms | Adj. p-value |
|---|---|
| MAP kinase tyrosine/serine/threonine phosphatase activity | 2.332e-4 |
| Calnexin/calreticulin cycle | 2.670e-3 |
| VEGFA-VEGFR2 Signaling | 3.575e-3 |
| RAF-independent MAPK1/3 activation | 6.532e-3 |
| Nectin adhesion pathway | 7.282e-3 |
| RHO GTPases activate IQGAPs | 8.947e-3 |
| Cadherin binding | 0.0107 |

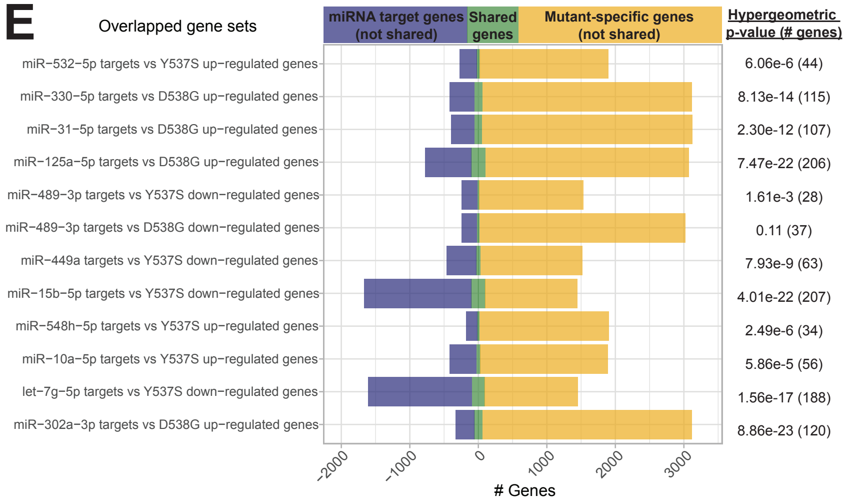


**Figure S1**. **miRNA target genes are enriched for mutant-specific genes.** Overlaps between Y537S, D538G, or both mutant-specific up- or down-regulated genes with predicted target genes for miR-301b (**A**), miR-181c (**B**), miR-331 (**C**), and let-7f (**D**) are shown. P-values indicating the significance of enrichment were calculated using the hypergeometric test. (**E**) Overlaps between Y537S or D538G up- or down-regulated genes and predicted miRNA target genes are shown for the remaining miRNA selected for further analysis. Blue bars indicate predicted miRNA target genes that are not mutant-specific genes, yellow indicates mutant-specific genes that are not predicted miRNA target genes, and green indicates genes that are both predicted miRNA target genes and mutant-specific genes. P-values indicating the significance of enrichment were calculated using the hypergeometric test.

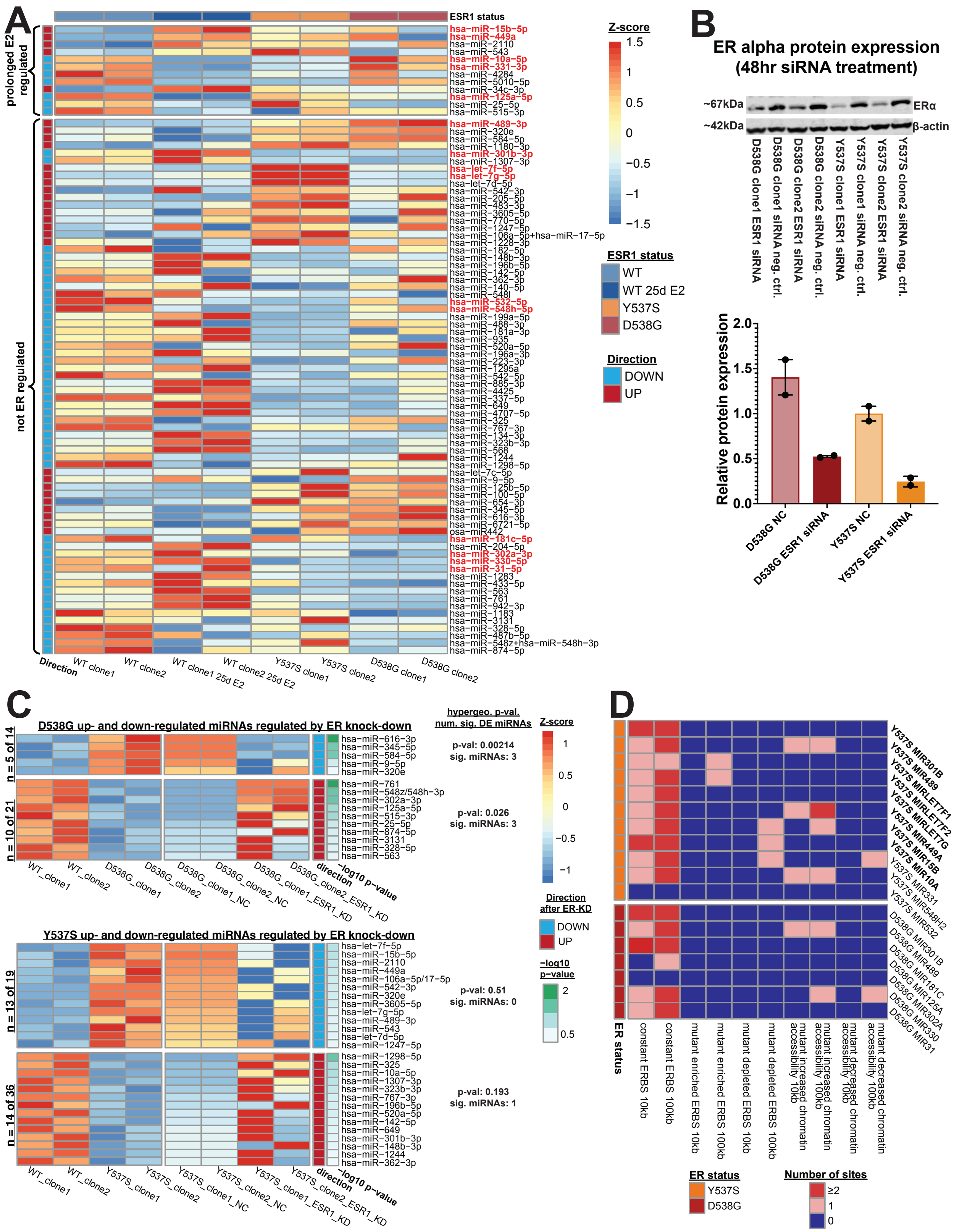


**Figure S2. ER contributes to altered miRNA expression.** (**A**) Heatmap shows expression levels for 84 miRNAs in WT clones without E2 treatment, WT clones treated with E2 for 25 days, Y537S clones without E2 treatment, and D538G clones without E2 treatment. 15 miRNAs selected for further investigation are highlighted in red. (**B**) Immunoblot analysis shows ER protein expression after siRNA treatment against *ESR1* in WT and ER mutant cells.

Quantification of immunoblot is shown with significance determined using a paired one-tailed Student's *t* test. (**C**) Heatmap shows expression levels for 42 of the 84 differentially expressed miRNAs that exhibit altered expression upon ER knock-down in mutant-ER cells in the expected direction. P-values for differential expression between ER knock-down and negative control treatments are shown in green and were calculated using a paired Student's *t* test. (**D**) Heatmap indicates the number of constant, mutant-enriched, or mutant-depleted ER binding sites or altered chromatin accessibility regions within 10kb or 100kb of the pri-miRNA transcription start site of differentially expressed miRNAs. Bolded miRNAs indicate miRNAs that were differentially expressed after ER knock-down.

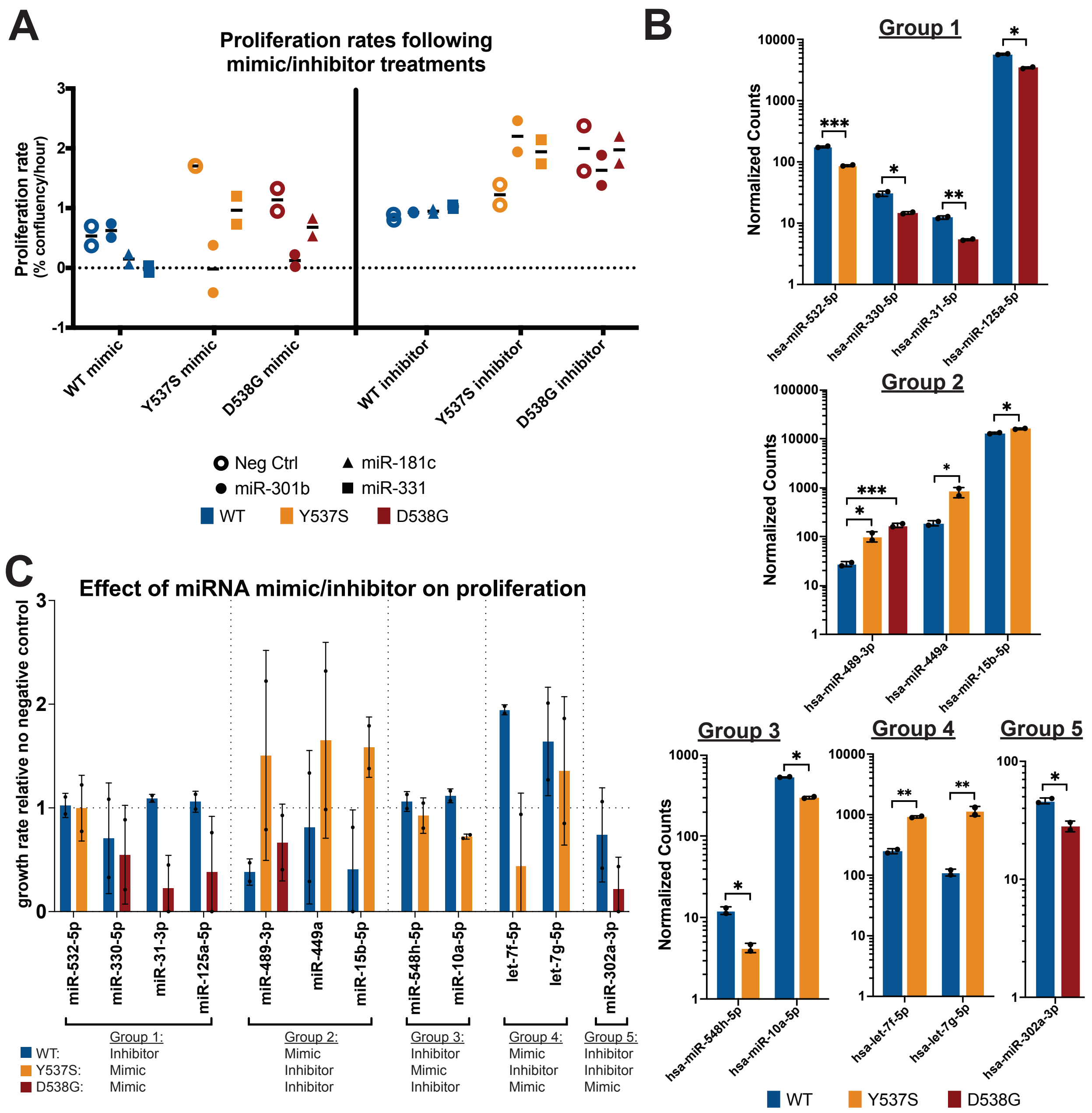


**Figure S3. Altered miRNA actvity impacts proliferation in ER mutant cells.** (**A**) Graph shows proliferation rates in percent confluency per hour for WT, D538G, and Y537S mutant cells treated with miRNA mimic or inhibitor for miR-301b, miR-331, miR-181c, or negative control. Line shows mean value of two replicate clones for each mutant ER genotype. (**B**) Expression levels are shown for 11 differentially expressed miRNAs in ER mutant cells. Normalized counts are shown on a log10 scale and miRNAs are organized into 5 groups based on their expression patterns and the mimic or inhibitor treatments used. Group 1 contains miRNAs that were down-regulated significantly or noticably in both ER mutants. Group 2 contains miRNAs that were significantly or noticably up-regulated in both ER mutants. Group 3 contains miRNAs that were down-regulated in the Y537S ER mutant cells only. Group 4 contained two miRNAs that were up-regulated in the Y537S ER mutant cells only. Group 5 contained a miRNA that was down-regulated in the D538G ER mutant only. Significance was determined using a Student's *t* test. (**C**) Growth rates relative to mimic or inhibitor negative control treatments are shown for each miRNA mimic or inhibitor treatment in WT (blue), Y537S (yellow), and D583G (red) cell lines.

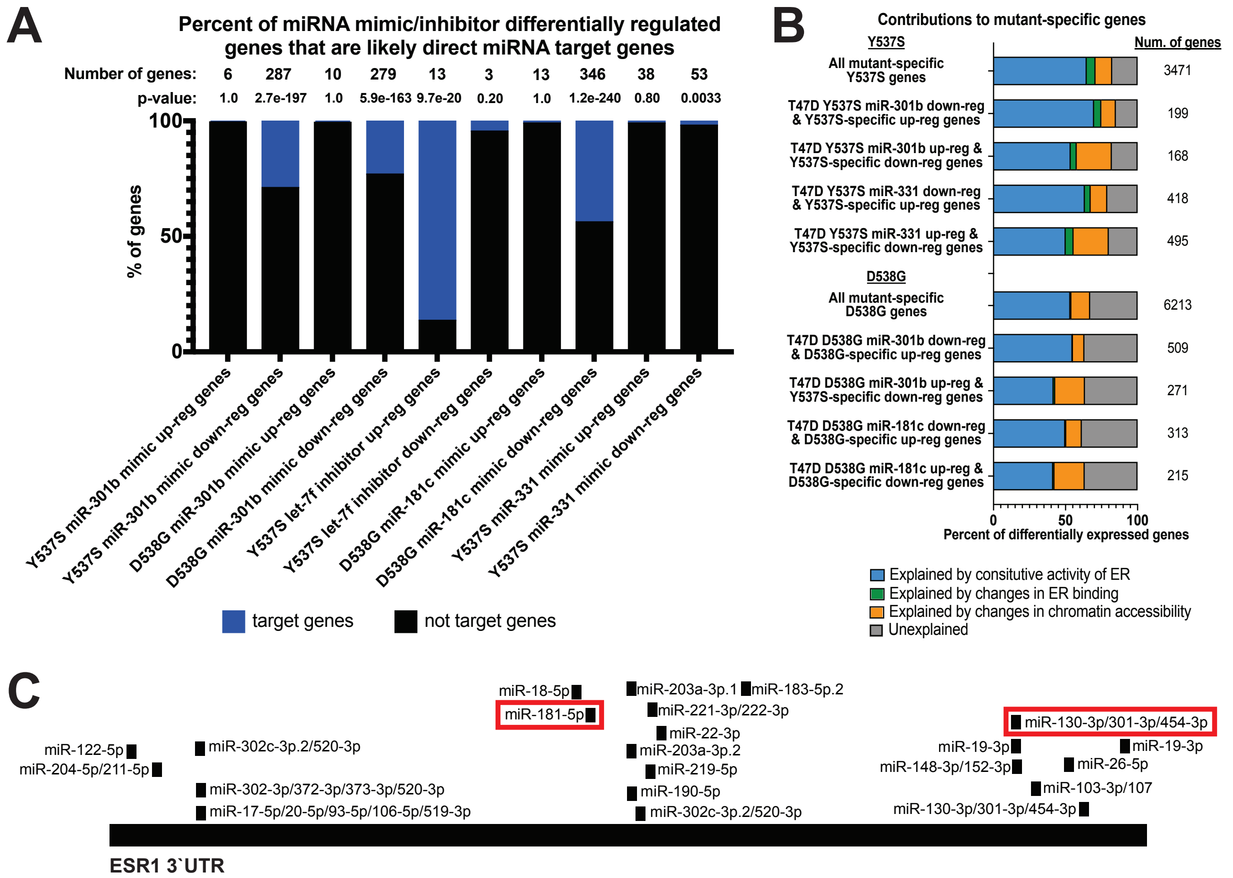


**Figure S4. miRNAs regulate mutant-specific genes.** (**A**) Bar graph shows the percent of miRNA mimic/inhibitor differentially regulated genes that are predicted direct targets for the corresponding miRNA. Hypergeometric tests were used to determine p-values. (**B**) miRNA-altered mutant-specific genes overlap with different classes of mutant-specific genes including those explained by mutant ER constitutive activity (blue), mutant ER driven altered chromatin accessibility (orange), mutant ER driven differential genomic binding (green), and genes that were previously unexplained (gray). Gene set is indicated on the left and the total number of genes for the corresponding gene set is listed on the right. (**C**) Figure represents *ESR1* 3'UTR and the regions targeted by miRNAs with the miR-301b and miR-181c target sites are boxed in red.

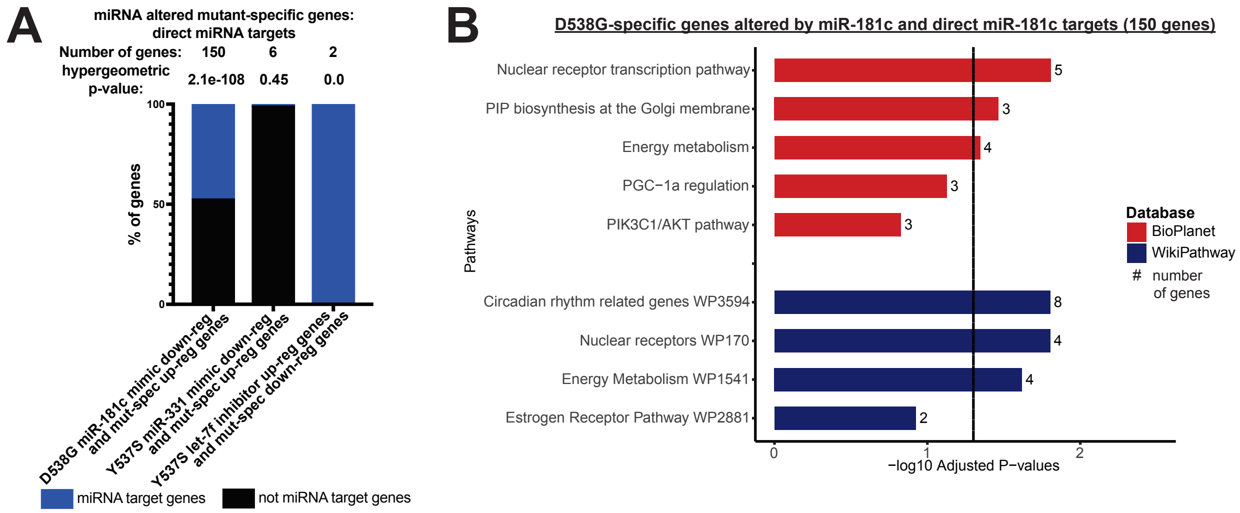


**Figure S5. miRNAs in ER mutant cells directly target mutant-specific genes involved in important cell signaling pathways.** (**A**) Percent of miR-181c, miR-331, or let-7f altered genes mutant-specific genes that are predicted direct targets of miR-181c, miR-331, or let-7f, respectively. Hypergeometric tests were used for p-value determination. (**B**) Bar graphs show pathways found to be enriched in miRNA-altered mutant-specific genes. -$\log_{10}$ adjusted p-values are indicated and the number of genes in each pathway gene set are shown.

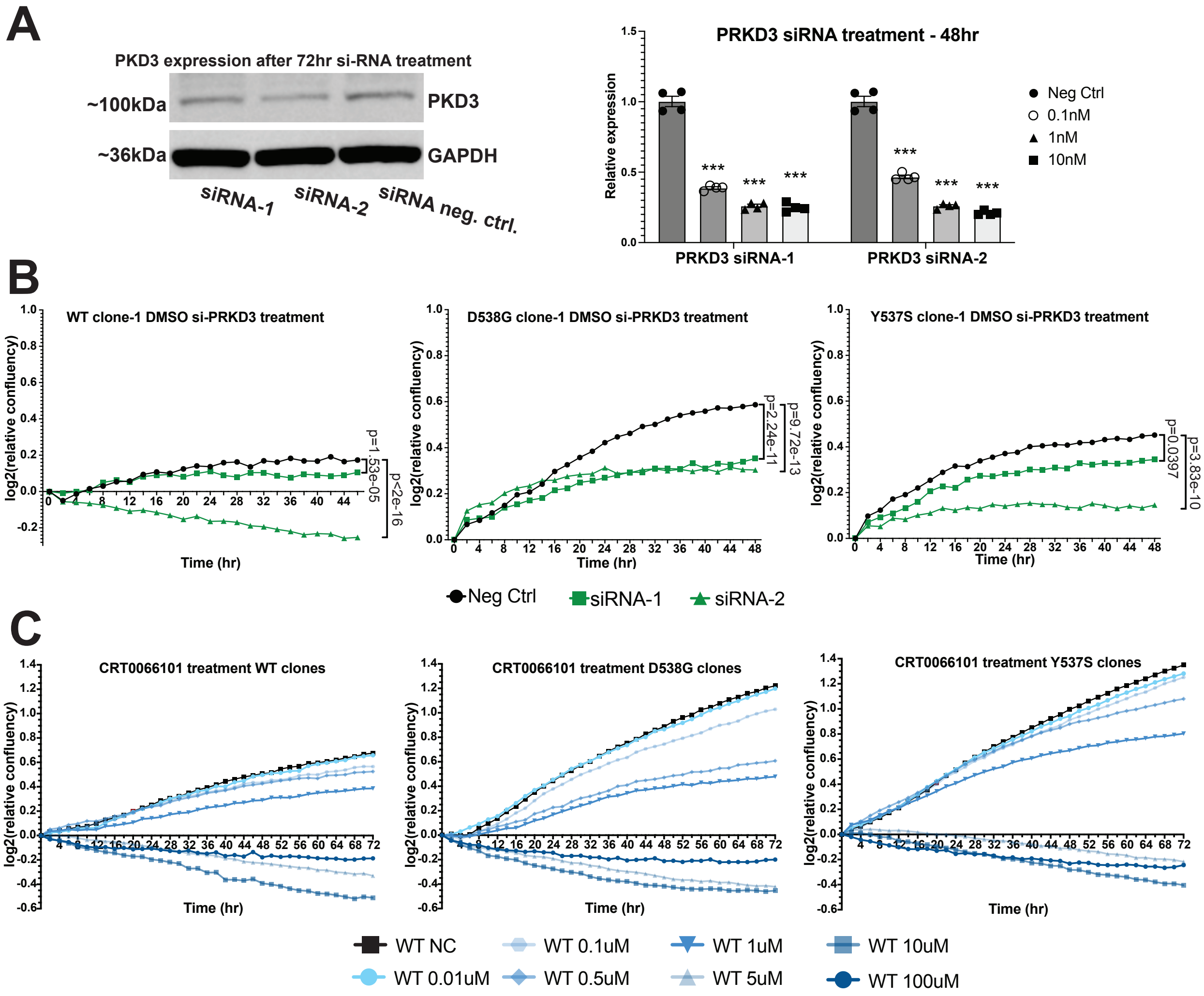


**Figure S6. Inhibition of *PRKD3* (PKD3) reduces proliferation in ER WT and mutant cells.** (**A**) Immunoblot analysis shows PKD3 protein expression after siRNA treatment against PRKD3 in WT ER cells. Bar graph shows qPCR analysis of *PRKD3* expression in T-47D cells after treatment with 0.1nM, 1nM, or 10nM siRNA against *PRKD3* or a negative control. Bars represent average ± SEM of four replicates for each treatment. (**B**) Line graphs show $log_2$ transformed relative confluence over time of WT or mutant ER cells treated with anti-*PRKD3* siRNAs or negative control. Significance between the slopes of the lines was determined using a generalized linear model and the Wald test. (**C**) Line graphs show $log_2$ transformed relative confluence over time of WT or mutant ER cells treated with pan-PKD inhibitor CRT0066101.

### SUPPLEMENTARY TABLES

**Supplemental Table S3. Percent overlap between mimic or inhibitor differentially expressed genes and all mutant-specific genes and unexplained mutant-specific genes**

| miRNA | mimic/inhibitor genes vs mutant-specific genes | % mimic or inhibitor genes | % mutant-specific genes | Hypergeometric p-value |
|---|---|---|---|---|
| miR-301b | Y537S mimic up-regulated vs Y537S mutant-specific down-regulated genes | 22.46% | 10.84% | 1.82e-53 |
| | Y537S mimic down-regulated vs Y537S mutant-specific up-regulated | 20.22% | 10.36% | 3.80e-41 |
| | D538G mimic up-regulated vs D538G mutant-specific down-regulated | 27.24% | 8.91% | 7.51e-44 |
| | D538G mimic down-regulated vs D538G mutant-specific up-regulated | 42.77% | 16.05% | 8.67e-167 |
| miR-181c | D538G mimic up-regulated vs D538G mutant-specific down-regulated | 21.61% | 7.07% | 1.78e-20 |
| | D538G mimic down-regulated vs D538G mutant-specific up-regulated | 39.87% | 8.87% | 6.14e-91 |
| let-7f | Y537S inhibitor up-regulated vs Y537S mutant-specific down-regulated | 13.33% | 0.13% | 0.22 |
| | Y537S inhibitor down-regulated vs Y537S mutant-specific up-regulated | 26.98% | 0.88% | 1.72e-6 |
| miR-331 | Y537S mimic up-regulated vs Y537S mutant-specific down-regulated | 17.25% | 31.94% | 3.54e-120 |
| | Y537S mimic down-regulated vs Y537S mutant-specific up-regulated | 17.15% | 21.76% | 4.63e-67 |
| **miRNA** | **mimic/inhibitor genes vs unexplained mutant-specific genes** | **% mimic or inhibitor genes** | **% mutant-specific genes** | **Hypergeometric p-value** |
| miR-301b | Y537S mimic up-regulated vs Y537S down-regulated genes | 4.01% | 10.91% | 3.42e-10 |
| | Y537S mimic down-regulated vs Y537S up-regulated | 3.05% | 8.75% | 1.58e-5 |

| | | | | |
|---|---|---|---|---|
| | D538G mimic up-regulated vs D538G down-regulated | 9.95% | 11.67% | 1.06e-23 |
| | D538G mimic down-regulated vs D538G up-regulated | 15.80% | 15.71% | 5.38e-53 |
| miR-181c | D538G mimic up-regulated vs D538G down-regulated | 7.94% | 9.32% | 1.46e-13 |
| | D538G mimic down-regulated vs D538G up-regulated | 15.41% | 10.11% | 8.25e-33 |
| let-7f | Y537S inhibitor up-regulated vs Y537S down-regulated | 0.0 | 0.0 | 1.0 |
| | Y537S inhibitor down-regulated vs Y537S up-regulated | 3.17% | 0.58% | 0.20 |
| miR-331 | Y537S mimic up-regulated vs Y537S down-regulated | 3.45% | 36.0% | 1.83e-28 |
| | Y537S mimic down-regulated vs Y537S up-regulated | 3.61% | 25.66% | 4.0e-19 |

**Supplemental Table S4. Sequence for qPCR analyses primers and siRNA knockdowns**

| Name | Sequence |
|---|---|
| PRKD3_qPCR_F | CTCACACGGGAGAGTGTTACC |
| PRKD3_qPCR_R | TTCATGTCATGGCGAAAGAGAA |
| TBP_qPCR_F | CCACTCACAGACTCTCACAAC |
| TBP_qPCR_R | CTGCGGTACAATCCCAGAACT |
| PRKD3_siRNA1_sense | rGrArArGrCrArArUrUrGrArUrCrUrGrArUrArArArCrArATC |
| PRKD3_siRNA1_antisense | rGrArUrUrGrUrUrArUrCrArGrArUrCrArArUrUrGrCrUrUrCrArC |
| PRKD3_siRNA2_sense | rArGrUrGrUrUrGrArCrArArArUrCrUrCrUrUrArGrUrCrATC |
| PRKD3_siRNA2_antisense | rGrArUrGrArCrUrArArGrArGrArUrUrUrGrUrCrArArCrArCrUrGrU |
